# RNA uridyl transferases TUT4/7 differentially regulate miRNA variants depending on the cancer cell-type

**DOI:** 10.1101/2021.07.12.451846

**Authors:** Ragini Medhi, Jonathan Price, Giulia Furlan, Beronia Gorges, Alexandra Sapetschnig, Eric A. Miska

## Abstract

The human terminal uridyl transferases TUT4 and TUT7 (TUT4/7) catalyse additions of uridines at the 3′ end of RNAs such as the precursors of the tumour suppressor miRNA let-7, upon recruitment by the oncoprotein LIN28A. Consequently, let-7 family miRNAs are downregulated. Disruption of this TUT4/7 activity inhibits tumorigenesis and hence targeting TUT4/7 can be a potential anti-cancer therapy. In this study, we investigate TUT4/7-mediated RNA regulation in two cancer cell lines by establishing catalytic knockout models. Upon TUT4/7 mutation, we observe a significant reduction in miRNA uridylation, which results in defects in cancer cell properties such as cell proliferation and migration. With the loss of TUT4/7-mediated miRNA uridylation, the uridylated miRNA variants are replaced by adenylated isomiRs. Changes in miRNA modification profiles are accompanied by deregulation of expression levels in specific cases. Unlike let-7s, most miRNAs do not depend on LIN28A for TUT4/7-mediated regulation. Additionally, we identify TUT4/7-regulated cell-type-specific miRNA clusters and deregulation in their corresponding mRNA targets. Expression levels of miR-200c-3p and miR-141-3p are regulated by TUT4/7 in a cancer cell type specific manner. Subsequently, BCL2 which is a well-established target of miR-200c is upregulated. Therefore, TUT4/7 loss triggers deregulation of miRNA-mRNA networks in a cell-type-specific manner. Understanding of the underlying biology of such cell-type-specific deregulation will be key when targeting TUT4/7 for cancer therapy.

**Graphical Abstract:** 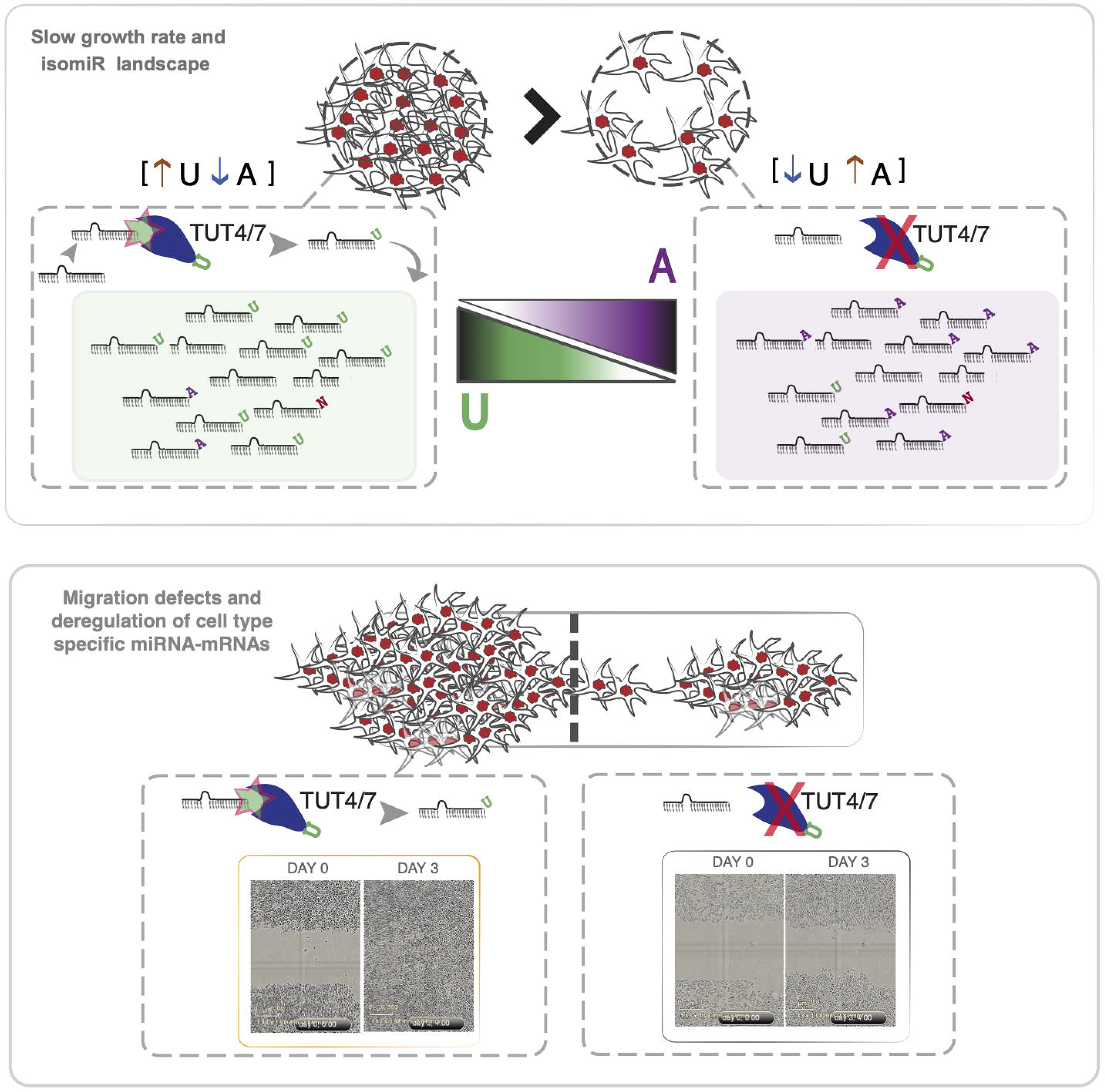

**Highlights:**

⍰ TUT4/7 catalyse 3′ uridylation of most miRNAs
⍰ Loss of uridylated isomiRs leads to the gain of adenylated isomiRs
⍰ TUT4/7-mediated miRNA deregulation and miRNA-mRNA interactions are specific to the cell line type
⍰ Loss of TUT4/7-mediated RNA uridylation inhibits cancer cell function

## INTRODUCTION

Endogenous small RNA populations include a distinct class of non-coding RNAs (ncRNAs) called microRNAs (miRNAs). miRNAs are well-characterized biomolecules with conserved roles in gene silencing. They are typically around 20-23 nucleotides (nt) in length with a seed region of 2-8 bases near the 5′ end that is able to pair perfectly or imperfectly with 3′ UTRs of target mRNAs (Bartel, 2004; Kim, Heo and Kim, 2010; Burroughs and Ando, 2019). Upon perfect pairing, miRNAs induce post-transcriptional silencing of the target via endonucleolytic cleavage, while imperfect pairing leads to translational repression (Bartel, 2004). Based on conservation of miRNA features (sequence, structure or function), various miRNAs have been classified under distinct families which are located in different genomic loci (Zou *et al*., 2014).

The miRNA loci are transcribed by RNA Pol II and processed at the 5′ and 3′ ends to generate capped and polyadenylated primary precursors of the miRNAs (pri-miRNAs). Subsequently, the RNase III enzyme DROSHA in conjunction with the RNA-binding partner DGCR8, measures ∼22 nt from the terminal loop of pri-miRNAs to introduce a cut and cleaves pri-miRNAs into ∼70 nt pre-miRNA precursors (Kim, 2005). Pre-miRNA hairpin precursors undergo further processing by another RNase III enzyme, DICER, to give rise to 5p and 3p mature miRNAs derived either from the 5′ arm or 3′ arm of the hairpin respectively. The mature miRNA derived from one arm of the pre-miRNA hairpin is preferentially loaded on a functional RISC (RNA-induced silencing complex) and the miRNA derived from the other arm is potentially degraded (Bartel, 2018). Additionally, the mature miRNA sequences of a specific miRNA gene is heterogeneous, with differences in 5′ end or 3′ end (additions or trimming), methylation, SNPs, and A-to-I editing (Barturen *et al*., 2014; Vitsios and Enright, 2015) being reported. Such miRNA variants are called isomiRs (Neilsen, Goodall and Bracken, 2012). Many isomiRs are miRNAs with templated additions (that align with reference sequence) or non-templated additions (3′ NTA) (Wyman *et al*., 2011).

With the advent of high-throughput sequencing, diverse 3′ NTA isomiRs have been identified for several miRNAs (Burroughs and Ando, 2019). Systematic analysis of the 3′ NTA profiles has revealed that such isomiRs are not experimental artefacts of sequencing techniques but are physiologically relevant (Wyman *et al*., 2011; Knouf, Wyman and Tewari, 2013). In addition, such non-templated additions are conserved across several species (Wyman *et al*., 2011; Saiselet *et al*., 2015). Overall, the most abundant modification irrespective of the studied system is addition of uridines (U) or adenines (A) (Chang *et al*., 2012; Li, Liao, *et al*., 2012; Saiselet *et al*., 2015). In the nematode *C. elegans*, uridylated miRNAs are higher in abundance than adenylated miRNAs. However, the reverse is true for higher mammals, such as humans and mice (Wyman *et al*., 2011). Some studies have highlighted that the 3′ NTA profiles change upon viral infection. For example, hepatitis B viral (HBV) infected human liver tissues have lower levels of adenylated miRNAs (Li, Tsai, *et al*., 2012). Another study has reported that B cells are enriched in adenylated miRNAs but uridylated isoforms dominate in their secreted exosomes (Koppers-Lalic *et al*., 2014).

In terms of proportions, mono-A or mono-U additions account for approximately 80 % of all NTAs, but the 3′ NTA landscape is not limited to these two. AA, AU, UA, and UU di-nucleotide terminal modifications are also observed and account for ∼1% of addition events (Li, Liao, *et al*., 2012; Li, Tsai, *et al*., 2012). Overall, the abundance of 3′ NTA isomiRs varies depending on the tissue-type or disease-context and even on the organism of study. Although, at a global level, the proportion of miRNA 3′ NTA is small, a subset of specific miRNAs has higher proportion (>75%) of isomiRs, which hints towards a biologically important role for their modified counterpart (Yang *et al*., 2019). Moreover, 3′ NTA additions have been shown to result in miRNA turnover and expression level changes with adenylation conferring stability in contrast to uridylation which promotes degradation (D’Ambrogio *et al*., 2012; Heo *et al*., 2012; Boseon Kim *et al*., 2015a; Gutiérrez-Vázquez *et al*., 2017).

In cancer, microRNA expression levels are useful in different aspects such as diagnosis, determining tumour progression, drug resistance and prediction of overall survival. For example, miR-200c and miR-141, both constituting the miR-200c/141 cluster, are diagnostic markers and are upregulated in ovarian cancer (Koutsaki *et al*., 2017; Sulaiman, Ab Mutalib and Jamal, 2016; Chen *et al*., 2019). These miRNAs mediate the various consequences ranging from increased cell proliferation and metastasis to drug resistance via their target mRNAs (Feng *et al*., 2014; Sulaiman, Ab Mutalib and Jamal, 2016). Therefore, it is crucial to understand how to modulate and control their expression levels to develop effective anti-cancer therapeutics.

The terminal RNA uridyl transferases TUT4 and TUT7 (TUT4/7) are the enzymes responsible for catalysing the addition of Us at the 3′ end of miRNAs such as the tumour suppressor miRNA let-7s and miR-324 to regulate their expression levels (Thornton *et al*., 2014). TUT4/7 mono-uridylate pre-let-7 miRNAs that have a 1 nucleotide overhang at their 3′ end, thereby restoring the optimal structure for efficient processing by DICER (Heo *et al*., 2012; B. Kim *et al*., 2015). However, in the presence of the oncogenic interactor LIN28A, TUT4/7 dwell longer at a pre-let-7 complex, oligo-uridylate it and mark it for decay (Yeom *et al*., 2011; B. Kim *et al*., 2015; Faehnle, Walleshauser and Joshua-Tor, 2017; Wang *et al*., 2017). In addition, TUT4/7 can uridylate pre-miR-324 (canonical with 2 nucleotide overhang), modulating the DICER cleavage site (Kim *et al*., 2020). In cancers such as glioblastomas overexpressing TUT4 and TUT7, pre-miR-324 is uridylated at the 3′ end, which leads to the 3′ arm derived miR-324-3p being more abundant than the miR-342-5p derived from the 5′ arm of the same precursor hairpin (Kim *et al*., 2020). Importantly, inhibition of TUT4/7-mediated uridylation results in a switch of the strand ratios with the miR-342-5p now being more abundant than the miR-324-3p (Kim *et al*., 2020). 5p/3p arm switching events have been observed in several independent studies and their ratios have been reported to change in tumour versus normal tissues for several cancer types (Li, Liao, *et al*., 2012; Li, Tsai, *et al*., 2012; Kim *et al*., 2020). In addition to arm switching, uridylation of miRNAs by TUT4/7 has implications for mRNA target selection. For instance, TUT4/7-mediated di-uridylation has been reported to enable the modified isomiRs to target non-canonical mRNA targets (imperfect seed pairing) via extensive base pairing at the 3′ end, thus expanding the miRNA target repertoire (Yang *et al*., 2019).

In this study, we explore the TUT4/7-mediated regulation of miRNA variants. We find that TUT4/7 catalyse most of the miRNA uridylation in the prostate cancer cell line DU145 and the ovarian cancer cell line IGROV1. We report that the loss of uridylated isomiRs observed in the TUT4/7 catalytic knockouts (TUT4/7 cKOs) results in a simultaneous gain of adenylated counterparts for specific miRNAs, and that this compensation is not a general phenomenon for all miRNAs. We present examples of miRNAs with no change in expression levels despite complete switch in isomiR populations. In addition, we identify cell line or cell type dependent and independent clusters of miRNAs that are deregulated upon TUT4/7 loss. Along with changes in isomiR population and miRNA abundances, we also observe overall slower growth rate and cell-type-specific defects in cell migration in the TUT4/7 cKOs. We also identify specific miRNAs such as miR-200c-3p and miR-141-3p to be regulated by TUT4/7 in a cell line specific manner. As miRNAs are increasingly being identified as cancer biomarkers and their expression and sequence profiles can distinguish between normal and tumour tissues (Lu *et al*., 2005), existence of such competing isomiR populations and TUT4/7-mediated cell type specific regulatory networks that control their expression levels hold clues to cancer progression. Thus, our findings provide a new avenue for the development of anti-cancer therapeutics as well as highlight the importance of further investigations for the cell type specific roles of TUT4/7 in cancer and the underlying isomiR biology.

## RESULTS

### The viable double knockouts of full-length TUT4 and TUT7 show cell type specific phenotypes

The RNA uridyl transferases TUT4 and TUT7 are broadly expressed in a diverse range of cell types. This study focuses on the prostate cancer cell line DU145 and the ovarian cancer cell line IGROV1, as both the cell lines have been previously reported to show reduced cell proliferation and tumour growth in xenograft models upon TUT4 transcript depletion via siRNAs or shRNAs (Piskounova *et al*., 2011; Fu *et al*., 2014). To understand the role of TUT4/7-mediated RNA uridylation in tumorigenesis, we generated double mutants of TUT4 and TUT7 in both cell lines by targeting exon 12 for *TUT4* and exon 14 for *TUT7* upstream of the catalytic domain encoding exon (exon 16 for both TUT4 and TUT7). Western blots probing for TUT4 and TUT7 show that both proteins are endogenously expressed at detectable levels in wildtype cells but are depleted in two independent double mutant clones of both cell lines (Figure 1A-1D), indicating that our CRISPR/Cas9 gene editing strategy was successful. Contrary to a previous study in cervical-derived HeLa cells, which concluded that the functional activity of both TUT4 and TUT7 is vital for cell viability (Lim *et al*., 2014), our data show that TUT4 and TUT7 are not required for viability in the prostate cancer cell line DU145 and the ovarian cancer cell line IGROV1. This suggests that the severity of cellular phenotypes upon TUT4/7 loss may be cell line or tumour specific.

**Figure 1.**
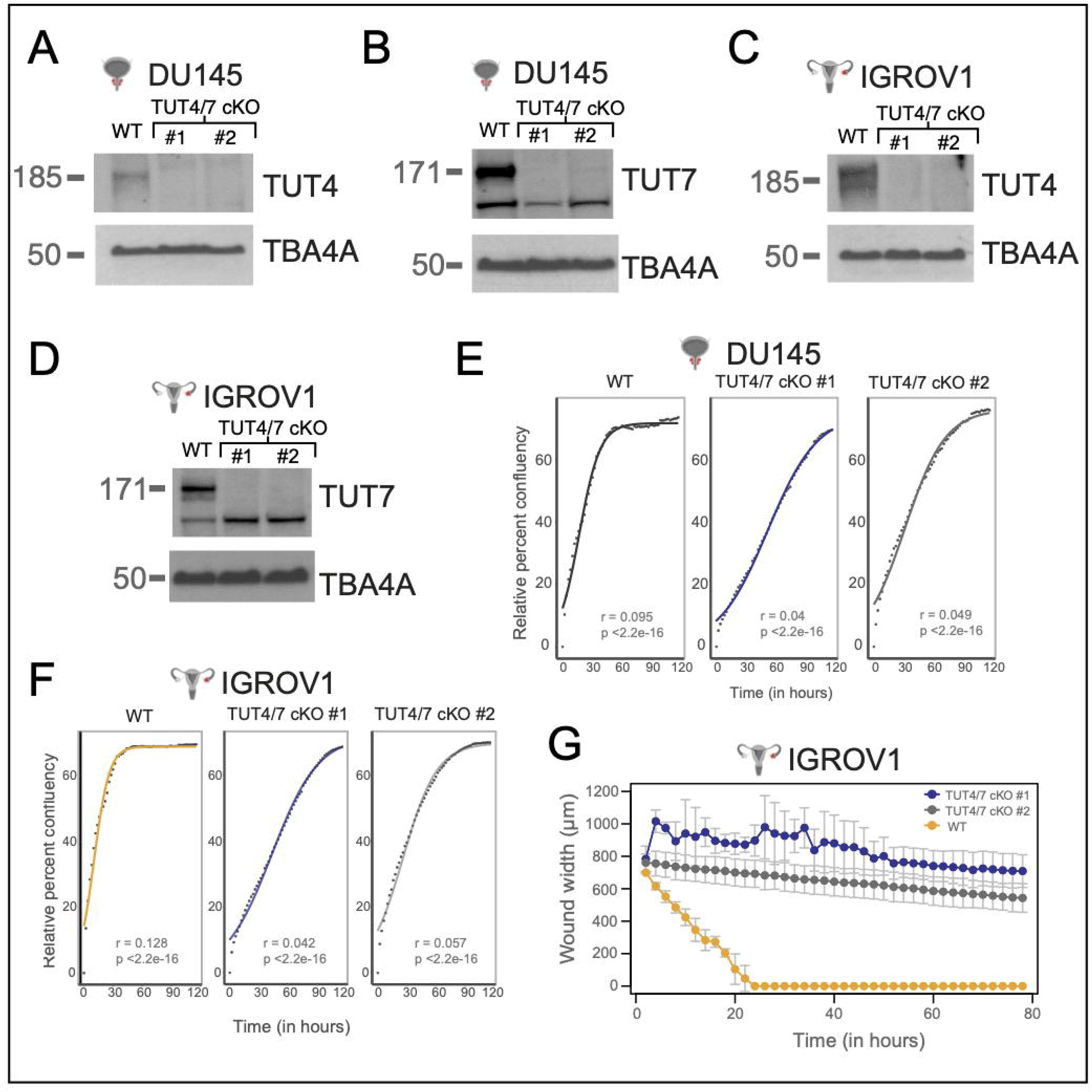
Loss of full-length TUT4 and TUT7 in the TUT4/7 double mutants has negative impact on cancer cell properties. Molecular masses of TUT4, TUT7 and the loading control a-tubulin (TBA4A) are 185 kDa, 171 kDa and 50 kDa, respectively. (A and B) Western blot images of TUT4 (A) and TUT7 (B) in independent TUT4/7 double mutant (cKO) clones (TUT4/7 cKO #1 and TUT4/7 cKO #2) of DU145 (n=3). (C and D) Western blot images of TUT4 (C) and TUT7 (D) in independent TUT4/7 double mutant (cKO) clones (TUT4/7 cKO #1 and TUT4/7 cKO #2) of IGROV1 (n=3). (E and F) Quantitative cell proliferation assay at 2-hour intervals using the Incucyte^®^ systems. Solid line indicates curve fitted to the logistic equation with r as the growth rate and p as the p-value of r. WT is the control for DU145 (E) and IGROV1 (F). TUT4/7 cKO #1 and TUT4/7 cKO #2 are the TUT4/7 double mutants. The y-axis represents relative percent confluency while the x-axis denote time in hours (n=2). Relative percent confluency at each time point was calculated by subtracting the initial confluency of seeded cells at the start point (time = 0) from the confluency at each time point. (G) Quantitative wound healing assays with measurements at 2-hour intervals using the Incucyt ^®^ systems. The y-axis denotes the wound width in μm and the x-axis represents time in hours. Cell line = IGROV1. WT is the control IGROV1 cell line. TUT4/7 cKO #1 and TUT4/7 cKO #2 are the TUT4/7 double mutants (n=2). See also Figure S1.

As the TUT4/7 double mutants of DU145 and IGROV1 were viable, we examined the TUT4/7 double mutants for possible defects in cancer cell properties such as cell proliferation, wound healing and cell migration. We scored for defects in cell proliferation using qualitative colony formation assays followed by crystal violet staining and quantitative cell proliferation assays for relative confluency using live-cell imaging at a 2-hour interval by the Incucyte systems. Defects in wound healing were also quantified using the Incucyte systems by live-cell imaging at regular intervals after the introduction of a scratch in a >90% confluent well. Impaired cell migration was assessed using trans-well migration assays followed by crystal violet staining. Compared to the wildtype cells, the TUT4/7 double mutants of DU145 and IGROV1 show a slower cell growth phenotype (Figure 1E-1F and Figure S1A-S1B). However, the IGROV1 double mutants of TUT4 and TUT7 also show a defect in wound healing unlike the DU145 TUT4/7 double mutants (Figure 1G and Figure S1C-S1D). Brightfield images at the start point (day 0) and end point (day 3) of scratch assays show that in the wildtype control the wound width is reduced to 0 at approximately the 20-hour timepoint, whereas the IGROV1 TUT4/7 double mutants fail to reform the cell monolayer at the end point of the assay (Figure 1G and Figure S1D).

To investigate whether the defect in wound healing is due to a cell migration defect and hence decoupled from the slow cell growth phenotype, we performed trans-well migration assays using 10% fetal bovine serum as a chemo-attractant. Both clones of DU145 TUT4/7 mutants do not show any drastic defect in migration (Figure S1E), while IGROV1 TUT4/7 mutants show slower cell migration compared to wildtype (Figure S1F). Note that cell migration is higher in the metastatic-site derived DU145 cancer cell line than the ovarian cancer cell line IGROV1 derived from stage III primary solid tumours (Figure S1E-S1F). Therefore, our results suggest that the loss of the full-length TUT4/7 have negative consequences on cancer cell properties and the severity may depend on the cancer cell line or the stage of cancer progression.

### Loss of full-length TUT4 and TUT7 results in a significant reduction of uridylated miRNAs

To assess the impact of TUT4/7 loss on miRNA populations, we performed small RNA sequencing and explored the modification profile of mature miRNAs. Using Chimira (version 2018; Vitsios and Enright, 2015), we first investigated the distribution of miRNA reads with terminal modifications (non-templated additions, i.e., NTA), A-to-I editing, SNPs or no modifications in wildtype cells and found that, overall, A-to-I editing and SNPs constituted a small fraction of the total population (<0.2%) (Figure S2A). NTA averaged at approximately ∼7% and ∼12% of the total miRNA reads for IGROV1 and DU145 respectively (Figure S2A). The majority of the reads (∼88% for DU145 and ∼93% for IGROV1) belonged to unmodified miRNA variants (Figure S2A), which contain canonical miRNAs (exactly matching the mature miRNA template), 3′ trimmed isomiRs (with only 5′ end matching to the ′ mature miRNA template), 3′ extended isomiRs (no 5′ variation but templated ′ extension in 3′ end), 5′ trimmed isomiRs (no 5′ variation), 5′ extended isomiRs (no 3′ variation but templated extension in 5′ end), and multi-length variants (both 5′ and 3′ end differs from the canonical mature miRNA) (Barturen *et al*., 2014).

Using sRNAbench (Barturen *et al*., 2014), we explored the distribution of the aforementioned isomiR variants in the two wildtype cell lines. Both canonical miRNAs (exact) and the 3′ extended isomiRs (lv3pE) constitute the highest fraction with each type occupying ∼36-38% of the total miRNA reads (Figure S2B). Approximately ∼8% of the total miRNA population comprises the 3′ trimmed isomiRs (lv3pT) (Figure S2B). 5′ extended isomiRs (lv5pE) constitute the lowest fraction (approximately ∼0.3-0.6%) of the unmodified pool (Figure S2B). This fraction is followed by 5′ trimmed isomiRs (lv5pT) and multi-length variants(mv) that constitute ∼2-4% of the total miRNA reads (Figure S2B). Reads with NTA fall in the remaining fraction (others) which is ∼10% and ∼13% of the total miRNA reads in IGROV1 and DU145 respectively (Figure S2B).

We found the 3p mature miRNAs to be more frequently modified than the 5p mature miRNAs in the two cell lines, potentially because of the pre-existing 3′ modifications of the precursor hairpins which also undergoes 3′ base additions before DICER processing. For example, TUT4/7 target pre-miRNAs such as pre-let-7s for terminal uridylation (Heo *et al*., 2012; B. Kim *et al*., 2015). Approximately ∼27% and ∼29% of the total number of miRNAs have 3p modified isomiRs in the DU145 and IGROV1 cell lines (Figure S2C). Both the cell lines share a high number of unmodified (overlap = 399) and modified (overlap = 167) 3p miRNAs (Figure S2D).

Next, we explored which one of the non-templated base additions (A, U, G or C) is predominantly observed on miRNA 3′ ends. On average, miRNAs with non-templated G and C additions constitute less than 0.4% of the total miRNA population (Figure 2A). Non-templated A additions represent the highest fraction of all base additions (∼8.6% for DU145 and ∼4.9% for IGROV1), followed by non-templated U additions or terminal uridylation (∼2.7% for DU145 and ∼4.2% for IGROV1) (Figure 2A).

**Figure 2.**
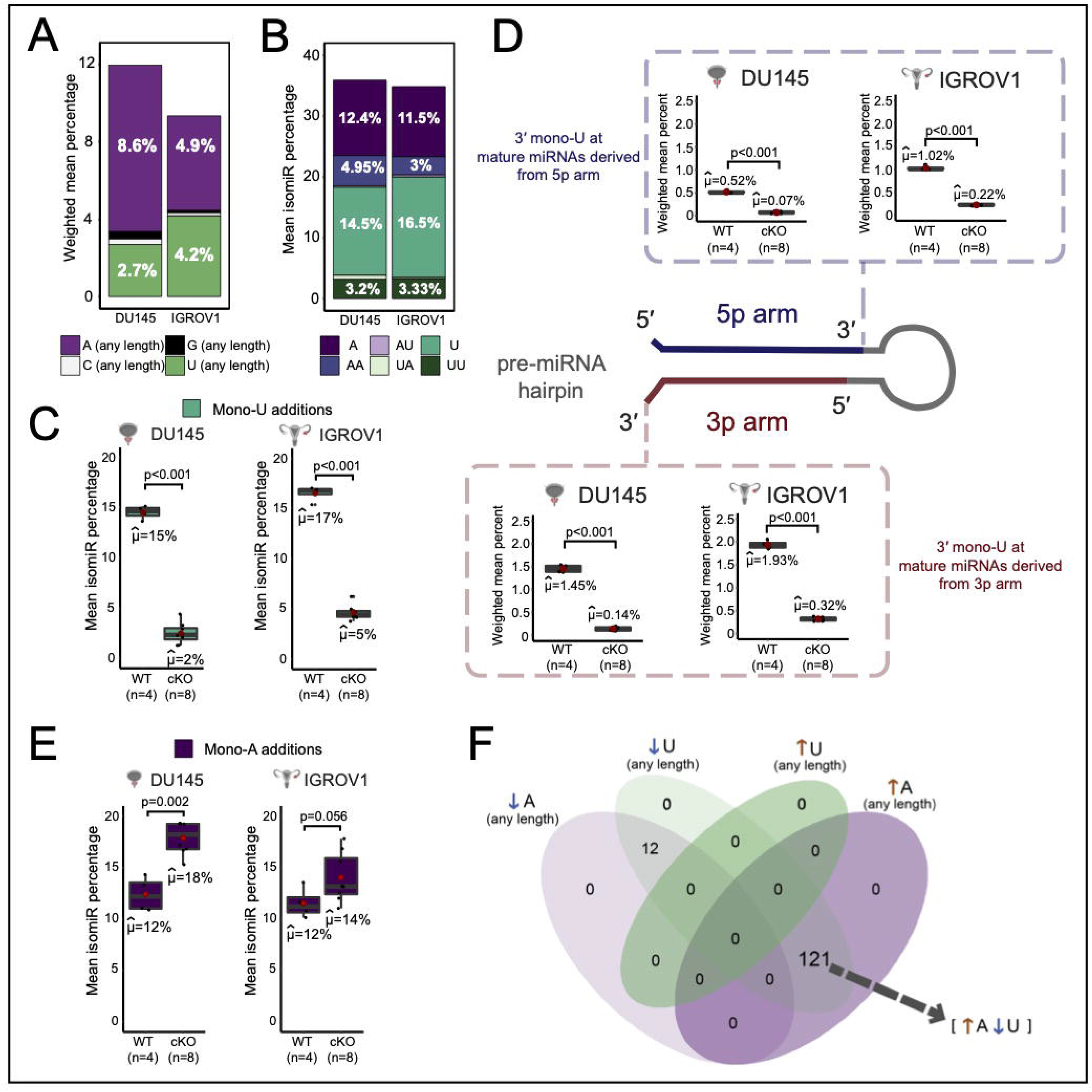
Impact of TUT4/7 loss on proportion of terminally modified miRNA variants. (A) “A (any length)”, “C (any length)”, “G (any length)”, and “U (any length)” are non-templated additions of A, C, G or U respectively with length 1 (in the case of mono additions) to the longest chain of homopolymeric tail detected at the end of a miRNA. (B) The isomiR ratio is the ratio of reads of a miRNA belonging to a particular NTA type to the total reads mapped to that miRNA. Mean isomiR percentage is the average isomiR ratio of the sample depicted in percentage. The NTA types displayed are mono-A additions (A), di-A additions (AA), mono-A addition followed by a mono-U addition (AU), mono-U additions (U), mono-U addition followed by a mono-A addition (UA) and di-U additions (UU). (C) Mono-U additions in the wildtype control (WT; n=4) and TUT4/7 double mutants (cKO; n=8) of DU145 and IGROV1. μ cap denotes the mean and p-value is indicated by p. (D) Mono-U additions at 3p and 5p mature miRNAs in the wildtype control (WT; n=4) and TUT4/7 double mutants (cKO; n=8) of DU145 and IGROV1. μ cap denotes the mean and p-value is indicated by p. (E) Mono-A additions in the wildtype control (WT; n=4) and TUT4/7 double mutants (cKO; n=8) of DU145 and IGROV1. μ cap denotes the mean and p-value is indicated by p. (F) Venn diagram showing overlap between miRNAs that show an overall decrease in adenylation (↓A), miRNAs with a decrease in uridylation (↓U), miRNAs with an increase in uridylation (↑U) and miRNAs with an increase in adenylation (↑A). See also Figure S2.

Within the non-templated base modifications, we investigated the mono-A (A), di-A (AA), mono-U (U), di-U (UU), AU and UA types of terminal additions as they were previously reported to be the most abundant 3′ base modifications (Li, Liao, *et al*.,′ 2012; Li, Tsai, *et al*., 2012). We found AU and UA types of NTA to be the least frequent, with a mean isomiR percentage below 1% (Figure 2B). Di-A (AA) and di-U (UU) types of NTA are observed at a frequency of ∼3-5% (Figure 2B). Single base additions of A and U are the most frequent types of non-templated additions (Figure 2B). Mono-uridylation displays a mean isomiR percent of 14.5% and 16.5% in both the DU145 and IGROV1 cell lines, a higher percentage than the one of mono-adenylation in the respective cell lines (Figure 2B). Therefore, our data show that mono-uridylation and mono-adenylation are the most frequent types of non-templated additions in the DU145 and IGROV1 cancer cell lines.

Loss of TUT4/7 results in a significant reduction of mono-U additions from isomiR percentages of 15% and 17% in the wildtype controls (WT) to 2% and 5% in the TUT4/7 DU145 and IGROV1 double mutants (Figure 2C). Although mono-U additions are one of the dominant forms of terminal uridylation, the abundance of U additions of any length at the 3′ end of mature miRNAs is reduced in the TUT4/7 double mutants (Figure S2E). The weighted mean ratio (ratio of number of reads with non-templated U to the total number of reads) has a significant reduction in the TUT4/7 double mutants of DU145 and IGROV1 compared to the wildtype (Figure S2E). Therefore, this validates that we have generated loss of function mutants of catalytic TUT4/7.

Next, we explored if TUT4/7 affect mono-U additions at the 3′ end of mature miRNAs derived from both the 3p and the 5p arm of the pre-miRNA hairpin and observed that the loss of TUT4/7 results in significant reduction of mono-U additions in the 3′ terminal end of both 3p and 5p mature miRNAs (Figure 2D). As 5p miRNAs are uridylated, TUT4/7-mediated uridylation occurs on mature miRNAs post DICER processing. However, mature miRNAs derived from the 3p arm are mono-uridylated at a higher frequency than the mature miRNAs derived from the 5p arm (Figure 2D). 3p mature miRNAs show a significant reduction in mono-uridylation from 1.45% and 1.93% of weighted mean in the wildtype controls to 0.14% and 0.32% in the TUT4/7 double mutants of DU145 and IGROV1 respectively (Figure 2D). In comparison to 3p mature miRNAs, 5p mature miRNAs are mono-uridylated at a lower mean frequency of 0.52% and 1.02% in the wildtype control which significantly reduces to 0.07% and 0.22% in the TUT4/7 double mutants of DU145 and IGROV1 respectively (Figure 2D). Therefore, mono-uridylation at the 3′ terminal end of 3p mature miRNAs is affected more than on 5p miRNAs upon TUT4/7 loss. However, overall terminal uridylation of any length at 3′ end of mature miRNAs decreases significantly in the TUT4/7 double mutants of both DU145 and IGROV1.

### Decrease in uridylated isomiRs leads to a simultaneous gain in adenylated counterparts

Previously, a few studies have noted an increase in the adenylated proportions of specific miRNAs upon the loss of TUT4/7. For example, adenylated reads for miR-324-3p simultaneously increase with loss of uridylated reads in TUT4/7 double knockouts of mouse cells (bone marrow, embryonic fibroblast, embryonic stem cell and liver) (Kim *et al*., 2020). In addition, ectopic expression of miR-27a-3p in TUT4/7 double knockouts and wildtype control of HEK293T show that the uridylated fraction of ectopically expressed miR-27a reduces by ∼2.5 fold with a concomitant ∼2.3 fold increase of adenylated reads for ectopic miR-27a-3p (Yang *et al*., 2019). Furthermore, mono-adenylation at 19 predicted targets of TUT4/7, which are all derived from 5p pre-miRNA arms, simultaneously increases upon TUT4/7 depletion and loss of mono-uridylation in the cervical cancer cell line HeLa (Thornton *et al*., 2014). In line with these previous observations, we observe a concurrent and significant global increase in mono-adenylation with the loss of mono-uridylation in the TUT4/7 double mutants of DU145 and IGROV1 (Figure 2E). The mean isomiR percentage of mono-adenylation increased from ∼12% in the wildtype controls to ∼18% and ∼14% in the TUT4/7 double mutants of DU145 and IGROV1, respectively (Figure 2E). This increase in terminal adenylation of mature miRNAs was not limited to the mono-adenylation type of non-templated A additions, but mature miRNA reads with A additions of any length increased in all the TUT4/7 double mutants (Figure S2F). The increase in adenylated isomiRs was higher in the DU145 TUT4/7 mutants compared to IGROV1 TUT4/7 mutants (Figure 2E). In contrast to mono-uridylation, mono-adenylation frequency is higher at terminal ends of 5p mature miRNA compared to 3p mature miRNA (Figure S2G). Mono-adenylation at 3′ end increased consistently and significantly for both 5p and 3p mature miRNA in the DU145 TUT4/7 mutants relative to the wildtype control (Figure S2G). However, this increase in mono-adenylated isomiRs was observed only in mature miRNA derived from the 3p arm in the TUT4/7 double mutants of IGROV1 compared to the wildtype (Figure S2G). Although there is an increase in mono-adenylation frequency, TUT2 enzyme which catalyses A additions and the exonuclease PARN which acts on miRNAs with A addition do not significantly deregulate at the RNA level (Figure S2H).

Out of the 167 overlapping modified 3p miRNAs between DU145 and IGROV1 wildtype cells (Figure S2D), 133 miRNAs had at least 1% terminal non-templated additions. In the TUT4/7 mutants of DU145 and IGROV1, we observed an increase in adenylated isomiRs coupled with a decrease in their uridylated counterparts in 121 miRNAs out of the 133 (Figure 2F). The miRNAs, let-7b-3p, let-7f-1-3p, let-7i-3p, miR-98-3p, miR-324-3p, miR-760, and miR-5001-3p gain adenylated isomiRs (miRNA reads with A-additions of any length) by at least 5-fold upon the loss of their uridylated counterparts (miRNA reads with U-additions of any length) (Figure 3A).

**Figure 3.**
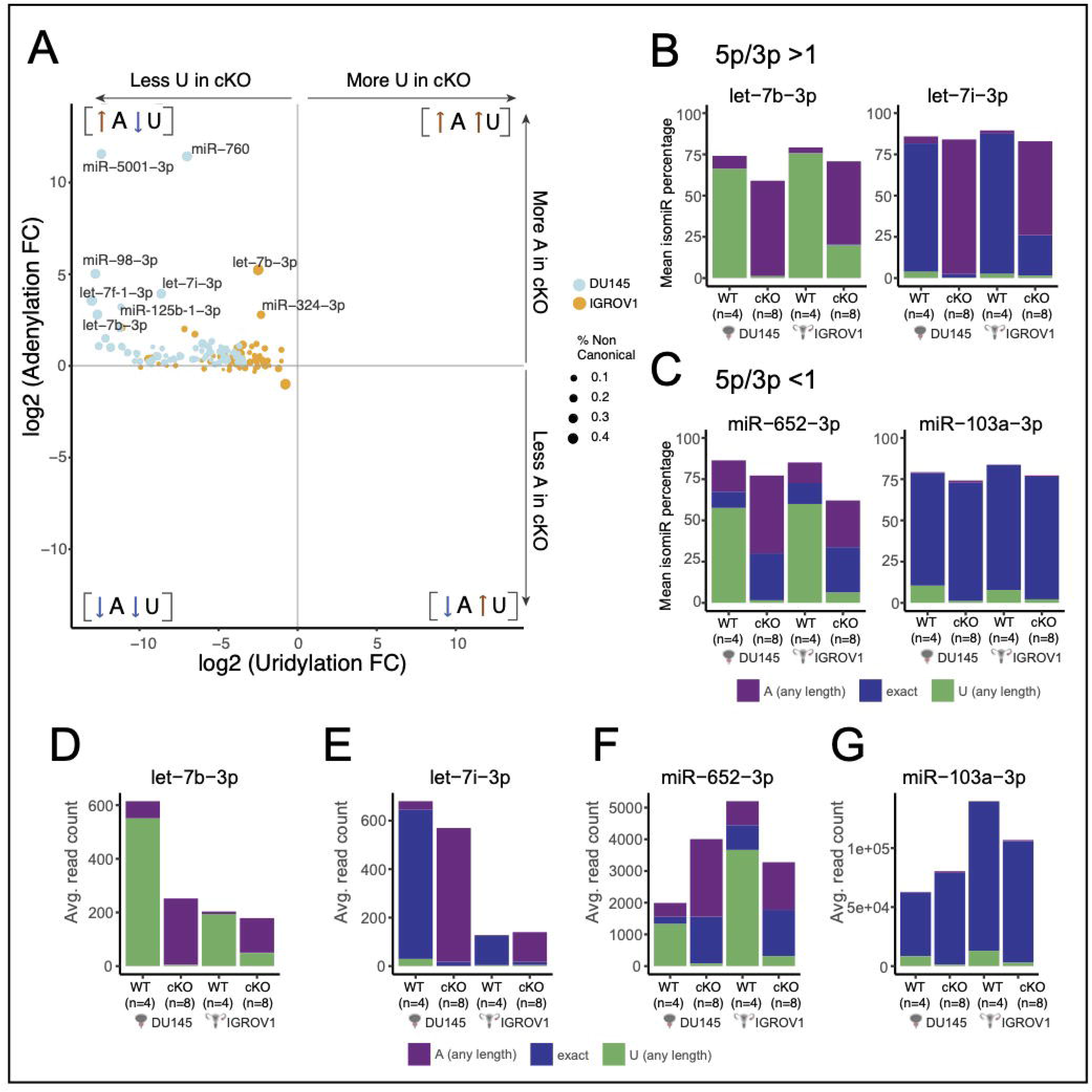
TUT4/7-dependent miRNAs gain adenylation upon loss of uridylation. (A) The 133 miRNAs out of the 167 overlapping modified miRNAs in DU145 and IGROV1 are depicted, and each miRNA has a terminal modification frequency of at least >1%. Fold change is calculated in the TUT4/7 double mutants compared to the wildtype control. The x-axis denotes logarithmic (base 2) fold-change in uridylation while the y-axis displays logarithmic (base 2) fold-change in adenylation in cKO versus WT. The miRNAs specific to IGROV1 are depicted in yellow and miRNAs specific to DU145 are displayed in blue. The labelled miRNAs are the ones with at least 5-fold increase in adenylation. (B) The y-axis indicates mean isomiR percentage. Both let-7b and let-7i have a higher 5p mature miRNA abundance compared to the 3p mature miRNA abundance, that is, a 5p/3p ratio of >1. Proportion of canonical, adenylated and uridylated isomiRs of let-7b-3p and let-7i-3p is displayed. (C) Proportion of canonical, adenylated and uridylated isomiRs of miR-652-3p and miR-103a-3p with 5p/3p<1 is displayed. (D, E, F and G) Abundances of 3p mature miRNAs of let-7b-3p (D), let-7i-3p (E), miR-652-3p (F), and miR-103a-2 (G) in the WT controls of DU145 and IGROV1 with their corresponding TUT4/7 double mutants (catalytic knockouts) represented by cKOs. See also Figure S3.

Next, we investigated if NTA modifications are preferentially added to either the 5p or 3p mature sequences of a specific miRNA to mark it for degradation. In our data, let-7b-5p and let-7i-5p are higher in abundance than let-7b-3p and let-7i-3p, respectively, with a ratio of 5p/3p>1. Both the 3p mature miRNAs of let-7b and let-7i have a higher proportion of modified isomiRs than the 5p miRNAs (Figure 3B, 3D-3E and Figure S3A-S3B). Although the canonical let-7i-3p with no terminal modification dominated the total miRNA pool in the wildtype control, the adenylated isomiRs emerged as the dominant isoform in the TUT4/7 double mutants of DU145 and IGROV1 (Figure 3B). Such upregulation of adenylated isomiRs upon the loss of uridylated counterparts is not limited to miRNAs with 5p/3p>1 but also observed for miRNAs with higher abundance of 3p miRNAs, i.e., 5p/3p<1. miR-652 with 5p/3p<1 also shows such changes in modified miRNA population (Figure 3C, 3F and Figure S3C). Therefore, presence of NTA might not be a mark for degradation for 5p/3p strand control. Examples of miRNAs that do not show an increase in adenylated isomiRs include miR-103a-2 (5p/3p<1). Uridylated miR-103a-3p is significantly reduced in the TUT4/7 catalytic knockouts but adenylated miR-103a-3p does not increase in abundance (Figure 3C, 3G and Figure S3D).

Similar to observations made in a recent study (Kim *et al*., 2020), the 3p mature miRNA of miR-324 was the dominant miRNA in the wildtype controls relative to the 5p mature miRNA and upon TUT4/7 loss, mature miRNAs derived from the 5p arm increased in abundance compared to the 3p mature miRNAs (Figure S3E). This arm switching mechanism is observed in both DU145 and IGROV1 TUT4/7 double mutants (Figure S3E). Thus, miR-324 displays a differential arm preference depending on TUT4/7 expression status. Both the 5p mature miRNA and 3p mature miRNA of miR-324 were marked by the simultaneous increase in adenylation upon the loss of TUT4/7 (Figure S3F). Such TUT4/7-mediated arm switching events were rare in our dataset and were not observed in most miRNAs. Figure S3G shows an example of a miRNA, miR-103a-2 which does not show arm switching. Thus, we conclude that loss of TUT4/7-mediated uridylation triggers a differential increase in terminal adenylation at the 3′ end for a subset of miRNAs.

### Specific miRNA clusters are deregulated upon the loss of terminal uridylation

3′ end uridylation and adenylation of RNAs have been associated with degradation and stability respectively (D’Ambrogio *et al*., 2012; Heo *et al*., 2012; B. Kim *et al*., 2015; Gutiérrez-Vázquez *et al*., 2017). TUT4/7-mediated uridylation has been shown to downregulate the expression levels of the let-7 family members when TUT4/7 interact with the stemness factor LIN28A (Lehrbach *et al*., 2009; Piskounova *et al*., 2011; Wang *et al*., 2017). Unlike IGROV1, DU145 cells do not express detectable LIN28A protein levels (Figure 4A). To explore the TUT4/7-mediated regulation of miRNA expression levels, we generated stable DU145 wildtype and DU145 TUT4/7 double mutant cell lines overexpressing lentiviral integrated CMV-driven LIN28A cDNA copies lacking introns and the 3’UTR (henceforth referred to LIN28A^OE^ DU145) and analysed their small RNA transcriptome (Figure 4A). A principal component analysis (PCA) shows that the first principal component (PC1) explains the majority (89%) of the variation in the dataset, which appears to be due to differences in the cell line type (Figure 4B), while the second principal component (PC2) separates IGROV1 TUT4/7 mutants from the wildtype controls (Figure 4B).

**Figure 4.**
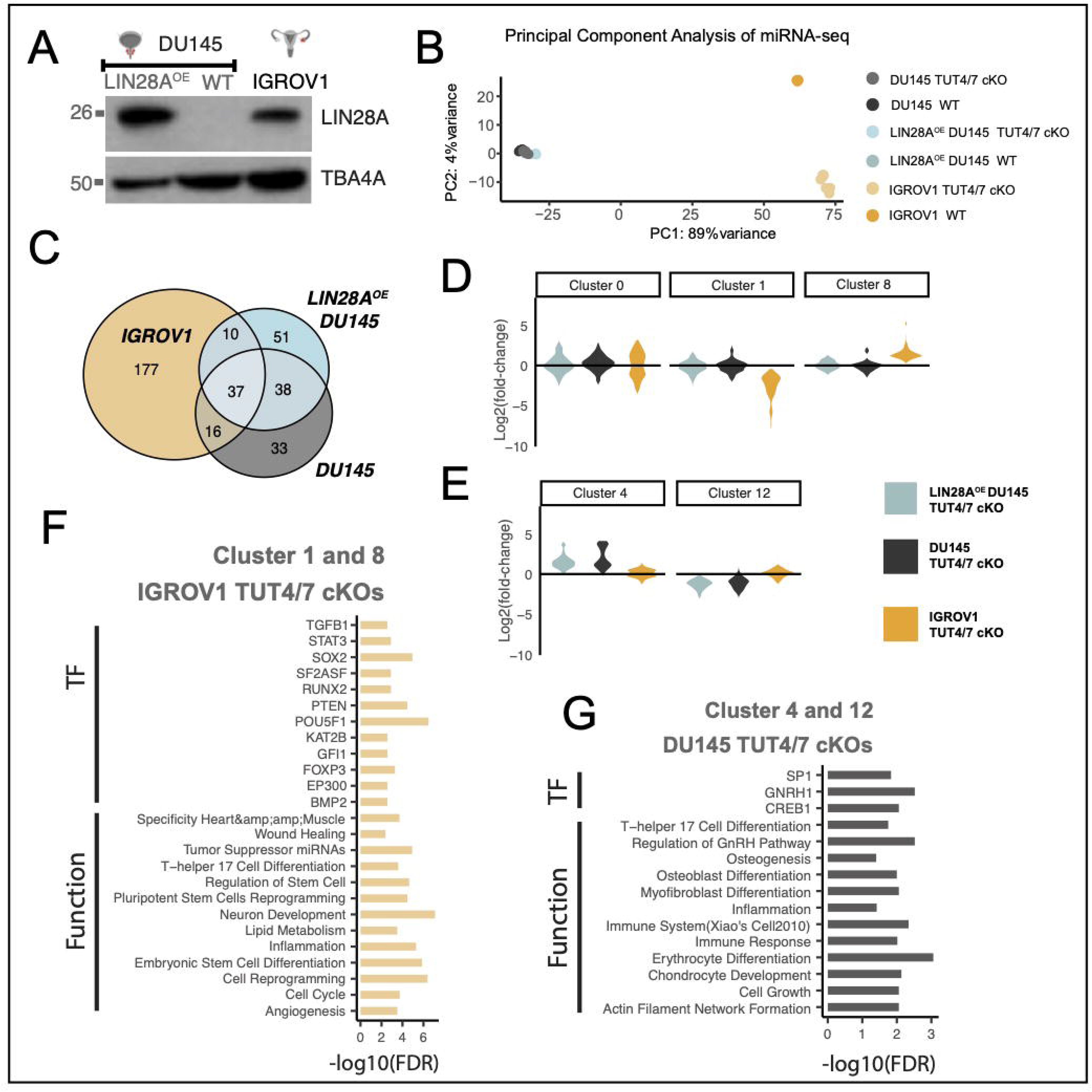
TUT4/7 loss results in cancer cell type specific miRNA deregulation. (A) Western blot image of LIN28A in the stable cell line LIN28A^OE^ DU145, DU145 (depicted as WT) and IGROV1 with a-tubulin (TBA4A) as a control (n=2). Molecular mass of LIN28A is 26 kDa and of TBA4A is 50 kDa. (B) Principal component analysis of miRNA dataset. PC1 versus PC2 in miRNA data for all genotypes. (C) Venn diagram showing gene overlap. (D and E) Violin plots displaying the expression trends in Clusters 0, 1 and 8 (D) and Clusters 4 and 12 (E). Fold change is calculated in the TUT4/7 double mutants (TUT4/7 cKO) relative to their respective wildtype control. (F and G) miRNA set enrichment analysis was performed with TAM 2.0 (Li et al., 2018). Enriched associations for the miRNA Cluster 1 and Cluster 8 with a similar deregulation pattern in TUT4/7 double mutants of IGROV1 (F) and for the miRNA Cluster 4 and Cluster 12 with a similar deregulation pattern in TUT4/7 double mutants of DU145 (G). See also Figure S4.

A PCA on all genotypes of the prostate cancer DU145-derived cell lines shows that PC1 versus PC2 captures clonal variation whereas PC2 vs PC3 captures the variation in the data that is associated with the presence or absence of TUT4/7 (Figure S4A). Of note, a higher number of miRNAs were deregulated in the IGROV1-derived mutants relative to the DU145-derived TUT4/7 catalytic knockouts (Figure 4C and Figure S4B). Therefore, the absence of TUT4/7-mediated miRNA regulation has a higher impact on the IGROV1 cell line relative to the prostate cancer cell line DU145. Next, we explored the effect of loss of uridylation on the expression levels of let-7 family members in a LIN28A-dependent context. We divided the let-7 family into the 3′ following classes: (i) canonical Group I miRNAs with 2 nucleotide overhangs at 3′ end (1), (ii) non-canonical Group II miRNAs with 1 nucleotide overhang at 3′ end (2), (iii) CSD+ let-7s which contains the (U)GAU binding motif for the cold shock domain (CSD) of LIN28A (+) and (iv) CSD-let-7s which do not contain the sequence motif for LIN28A binding via its CSD domain (-).

In DU145, all let-7 miRNAs are generally either downregulated or unchanged in TUT4/7 cKOs (Figure S4C). Based on previous data (Ustianenko *et al*., 2018), miRNA from CSD+ precursors should be upregulated in LIN28A-positive TUT4/7 cKOs. This trend is true for all CSD+ let-7 miRNAs except miR-98 in DU145-derived TUT4/7 cKOs (LIN28A positive vs LIN28A negative) (Figure S4C). However, we do not observe the same trend of upregulation for CSD+ let-7s in IGROV1 TUT4/7 catalytic knockouts which endogenously express LIN28A (Figure S4C). Hence, this suggests that TUT4/7-mediated let-7 regulation is cell line-specific and does not always depend on LIN28A in a cellular environment, contrary what is known from *in vitro* tailing reactions (Piskounova *et al*., 2011; Thornton *et al*., 2012; B. Kim *et al*., 2015).

Next, to identify miRNAs that depend on LIN28A-TUT4/7 for the regulation of their expression, we performed k-means clustering (k=15) of all miRNAs excluding let-7 family members and identified 14 miRNA clusters (Figure S4D). Cluster 0 (number of miRNAs, N=60) consists of miRNAs whose expression does not significantly change (Figure 4D). Cluster 1 contains miRNAs (N=76) that are downregulated and Cluster 8 miRNAs (N=34) that are upregulated only in the IGROV1 TUT4/7 cKOs (Figure 4D). Similarly, Cluster 4 represents miRNAs (N=32) that are upregulated and Cluster 12 miRNAs (N=9) that are downregulated only in the DU145-derived TUT4/7 cKOs (Figure 4E). On performing miRNA set enrichment analysis using TAM 2.0 (Li et al., 2018), we found that only IGROV1 TUT4/7 cKO-specific clusters (Cluster 1 and 8) display “wound healing” as an enriched function (Figure 4F), suggesting that deregulation of the miRNAs in these two clusters results in defective wound healing in the IGROV1 TUT4/7 double mutants. The miRNA set enrichment analysis for Cluster 4 and 12 show “cell growth” and “differentiation” related terms, suggesting that these miRNAs might contribute to the slow growth phenotype in the TUT4/7 double mutants of the metastatic-site derived DU145 cancer cell line (Figure 4G). In addition, miRNAs belonging to Cluster 2 (N=19) and Cluster 3 (N=81) show a general trend of downregulation and upregulation respectively independent of the cell line, with “differentiation” emerging as the most enriched function (Figure S4E-S4F). Therefore, TUT4/7-mediated regulation of miRNA expression levels is miRNA-specific, and subsets of miRNAs are regulated in a cell line specific manner.

### Cell type-specific negative correlation of miRNA-mRNA interactions upon functional loss of TUT4/7 activity

miRNAs mediate gene silencing by targeting mRNAs post-transcriptionally. Hence, global deregulation of miRNA abundance may severely impact the mRNA transcriptome. In order to assess the effects of TUT4/7 loss on the transcriptome, we sequenced poly(A)-selected transcripts in our samples. PCA analyses on our mRNA-seq datasets were similar to those performed on our miRNA datasets (Figure S5A-S5C). According to the current model of TUT4/7-mediated mRNA regulation (Lim *et al*., 2014), TUT4/7 are considered general mRNA decay factors and global upregulation of mRNA is expected upon the loss of uridylation. However, we did not observe this global trend of upregulation in the DU145 and IGROV1 TUT4/7 cKOs (Figure S5D). In addition, minimal overlap of deregulated mRNAs between the cell lines suggests that TUT4/7 differentially regulate mRNAs depending on the cell line (Figure S5E). Upon gene-ontology analysis of the deregulated mRNAs, “immune response” pathways were enriched in the prostate DU145 TUT4/7 cKOs (Figure S5F). This is not unusual given the role of TUT4/7 in anti-viral processes and transposable element regulation (Le Pen *et al*., 2018; Warkocki *et al*., 2018). In comparison, a higher number of processes relating to “wound healing” and “cell migration” were enriched in the ovarian IGROV1 TUT4/7 cKOs (Figure S5F-S5H).

Unlike the DU145 cell line derived from a metastatic site, the IGROV1 cell line is originally derived from a primary tumour in Stage III which is before the metastatic Stage IV (Stone *et al*., 1978; Bénard *et al*., 1985). During metastasis, the metastasis-associated miR-200c cluster are upregulated and it is also used as diagnostic markers in ovarian cancer (Sulaiman, Ab Mutalib and Jamal, 2016; Chen *et al*., 2019). We found that miR-200c-3p and miR-141-3p belonging to the miR-200c cluster are targets of TUT4/7 and are regulated in a cell line specific manner (Figure 5A-5D, Figure S6A-S6B and Figure S6D-S6G). Both miRNAs are repressed only in the IGROV1 TUT4/7 cKOs (Figure 5A-5D, Figure S6A-S6B and Figure S6D-S6G). In turn, the well-characterised target of miR-200c, the anti-apoptotic factor BCL2 (Feng *et al*., 2014) is now upregulated (Figure 5E-F, Figure S6C and Figure S6H-S6I). Although BCL2 has elevated expression levels in many cancers and is an anti-cancer target, overexpression of BCL2 alone in cells does not impart tumorigenic potential but rather can have pro-survival benefits (Lu *et al*., 1995; Kirkin, Joos and Zörnig, 2004; Czabotar *et al*., 2014).

**Figure 5.**
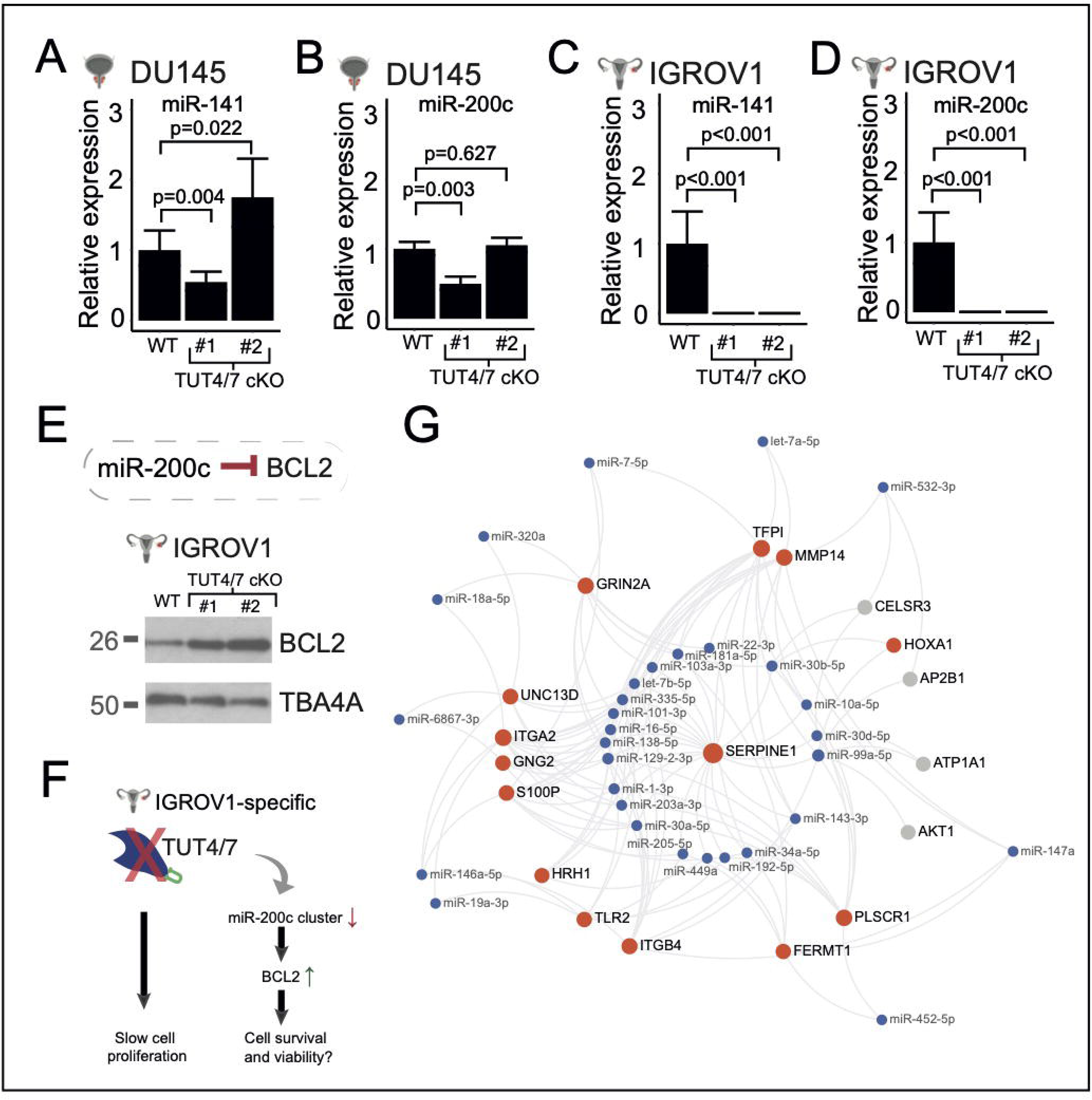
IGROV1-specific miRNA-mRNA interactions. (A, B) RT-qPCR quantification of miR-141-3p (A) and miR-200c-3p (B) in DU145 and derived TUT4/7 cKOs. Number of biological replicates (n) for WT = 4. TUT4/7 cKO #1 represents one independent clone (n=4) and TUT4/7 cKO #2 (n=3) represents the other. P-values are denoted by p and is determined by Student’s t-Test. (C, D) RT-qPCR quantification of miR-141-3p (C) and miR-200c-3p (D) in IGROV1 and derived TUT4/7 cKOs. Number of biological replicates (n) for WT = 4. TUT4/7 cKO #1 represents one independent clone (n=4) and TUT4/7 cKO #2 (n=4) represents the other. P-values are denoted by p and is determined by Student’s t-Test. (E) Downregulation of miR-200c in TUT4/7 cKOs is accompanied by upregulation of its target BCL2. (F) TUT4/7 loss results in IGROV1-specific downregulation of the miR-200c cluster, which in turn leads to upregulation of its target BCL2. Loss of TUT4/7 catalytic function results in slow cell proliferation. BCL2 might contribute to cell viability and survival of IGROV1 TUT4/7 catalytic knockouts. (G) Each connection from the miRNA in blue is to mRNA target that are implicated in cell migration or wound healing. Red dots denote mRNAs that are downregulated otherwise the mRNA is denoted in gray. This network was created using miRNet (Fan et al., 2016). See also Figure S5 and Figure S6.

Additionally, we used miRNet 2.0 (Chang *et al*., 2020) in order to identify miRNA-mRNA interactions related to wound healing and cell migration in the IGROV1-derived TUT4/7 cKOs. We used the list of IGROV1-specific downregulated mRNAs and upregulated miRNAs as query and gene ontology terms related to wound healing or cell migration were selected. Subsequently, the miRNA-mRNA interaction network in Figure 5G was obtained. Many cancer-cell invasion and migration promoting genes such as SERPINE1, PLSCR1, ITGA2, MMP14, etc. are significantly downregulated in the IGROV1 (Figure 5G). According to our network, these mRNAs interact with upregulated miRNAs such as those belonging to the miR-30 family (miR-30a-5p and miR-30b-5p) and miR-99a-5p (Figure 5G). Although such negative correlation of expression levels of miRNA and their mRNA targets is widely observed in this study, we also observe several examples where a negative correlation is not observed. For example, despite the trend of downregulation in their mRNA targets (Figure 5G), let-7a-5p and let-7b-5p expression levels do not drastically change. In addition, both miR-7-5p and its targets are downregulated in the IGROV1 TUT4/7 cKOs, thus showing a positive correlation with its targets. Such observations could be due to multiple miRNAs having shared targets.

In summary, TUT4/7 are master regulators and versatile in their RNA regulation function. TUT4/7-mediated miRNA regulation depends on the cell line of study. This leads us to hypothesise that certain, still unknown, cell type specific upstream factors exert overall influence over TUT4/7 recruitment. This will require further investigations as our data suggests that the severity of TUT4/7 inhibition may differ depending on the cancer cell type or the stage of cancer progression. Thus, a better understanding of cell type specific interactors of TUT4/7 will be crucial in stratifying tumours for potential treatment with TUT4/7 inhibitors.

## DISCUSSION

TUT4/7-mediated RNA metabolism is complex and shapes the transcriptomic network in a cell-type-dependent manner. Here, we explored the global TUT4/7-mediated RNA regulatory networks in two different cancer cell line models with distinct tissues of origin. The two cell lines used are the prostate cancer cell line DU145, which is derived from a metastatic site and has a high stem cell-like population and the cell line IGROV1 derived from the primary solid tumour of stage III ovarian cancer (Stone *et al*., 1978; Bénard *et al*., 1985; Pfeiffer and Schalken, 2010). We found that TUT4/7 catalytic knockouts display an overall slower cell proliferation, but with cell line specific differences in cell migration phenotype (Figure 1E-1G and Figure S1A-S1F). Such cell line specific phenotypic defects could be due to loss of TUT4/7-mediated 3′ uridylation of specific miRNAs (Figure 2C and Figure S2E). Based on k-means clustering, the deregulated miRNAs can be divided into groups which follow the same trend of deregulation in the TUT4/7 catalytic knockouts (Figure S4D). These groups can be further classified into distinct miRNA clusters that are either cell type-independent or -dependent (Figure 4D-4E and Figure S4E). TUT4/7 loss has a greater impact on the cancer cell properties of the ovarian cancer cell line IGROV1 than the prostate cancer cell line DU145. GO terms relating to cell migration are exclusively limited to the IGROV1-specific miRNA clusters whereas terms relating to differentiation emerge in the DU145-specific clusters (Figure 4F-4G and Figure S4F). In addition, we identify deregulated mRNA targets relating to cell migration exclusively in the ovarian cancer cell line (Figure S5F-H). The miRNAs miR-200c-3p and miR-141-3p are regulated by TUT4/7 in a cell type dependent manner (Figure 5A-5D and Figure S6A-S6G). Therefore, our results suggest that TUT4/7-dependent miRNA-mediated mRNA regulation varies depending on the cancer cell line of study, which results in observed differences in defects of cancer cell properties. These inherent differences in TUT4/7-mediated RNA regulation could also possibly depend on the cancer status and the stage of cancer progression, where TUT4/7 modulate transcriptomic changes based on the current requirements for tumorigenesis. Our data suggests that in DU145, TUT4/7-mediated miRNA regulation may help in maintaining an enriched population of stem cell-like cells as the removal of TUT4/7 function results in the emergence of “differentiation” related GO terms (Figure 4G). In addition, for the IGROV1 cell line, TUT4/7-mediated regulation may prepare the tumour for the next stage of cancer, i.e., metastasis, as cell migration, “wound healing” and “angiogenesis” related genes are deregulated upon TUT4/7 loss (Figure 4F and Figure S5G).

Moreover, we find that the regulation of the tumour suppressor let-7 family members, which is known to be dependent on the LIN28A-TUT4/7 pathway, does not strictly adhere to the previously postulated uniform downregulation model inside cells, and that LIN28A-TUT4/7-mediated regulation depends on the cell line of study (Figure S4C). Similar to observations made in a recent study (Ustianenko *et al*., 2018), this is contrary to the uniform downregulation model observed *in vitro* (Heo *et al*., 2008, 2009; Viswanathan, Daley and Gregory, 2008). Moreover, we find that, beyond let-7, miRNAs do not generally show a trend in expression level change that depends on the LIN28A-TUT4/7 pathway (Figure S4D).

Furthermore, loss of catalytic activity of TUT4/7 leads to an increase in adenylated miRNAs. This has been observed previously upon the depletion of TUT4/7 but was limited to few miRNA examples (Thornton *et al*., 2014; Yang *et al*., 2019; Kim *et al*., 2020). Here we provide a global view of changes in miRNA variant populations in the TUT4/7 double mutants of the prostate cancer cell line DU145 and the ovarian cancer cell line IGROV1 (Figure 2A-F and Figure S2A-G). Many effector enzymes similar to TUT4/7 such as TUT2, TENT4A and TENT4B have been identified and shown to catalyse A additions. Upon the loss of TUT4/7, as specific miRNAs undergo increased adenylation, it is possible that adenylating enzymes such as TUT2 and TENT4A/B are in competition with TUT4/7 for the same miRNA substrate. Therefore, in the absence of TUT4/7, the adenylating enzymes can now access these specific miRNAs and target them for adenylation as we observe an increase in adenylation of TUT4/7-dependent miRNAs in our data (Figure 2E and Figure S2F-G).

Additionally, reads for isomiRs with non-templated base additions were considered to be artefacts of sequencing technologies but this notion was subsequently dismissed and invalidated (Wyman *et al*., 2011; Knouf, Wyman and Tewari, 2013). One of the critical factors that proves their relevance in biological processes is the association of NTA modified isomiRs with active components of RNA-induced silencing complexes (RISCs) (Burroughs *et al*., 2010; Yang *et al*., 2019). Argonaute (AGO) proteins are responsible for loading miRNAs into RISC complexes. Existence of multiple AGO proteins may indicate selective and preferential sorting of miRNAs into individual AGOs. Indeed, miRNAs terminally modified with adenine additions have reduced association with AGO2 and AGO3 (Burroughs *et al*., 2010). High base complementarity between miRNA:mRNA is a factor for intrinsic endonuclease activity of AGO2, and adenine additions might impact its slicer activity (Liu *et al*., 2004; Song *et al*., 2004; Burroughs *et al*., 2010). However, other terminal base modifications did not show any significant changes in AGO-associations (Burroughs *et al*., 2010). Therefore, change in isomiR population for specific miRNAs in the TUT4/7 catalytic knockouts might also affect preferential AGO loading and the underlying consequences might need further investigation.

Another two important aspects of 3′ modified isomiRs are miRNA turnover and target regulation. Terminal oligo-uridylation of miRNAs has been associated with instability, whereas adenylation is attributed to increased stability of miRNAs (D’Ambrogio *et al*., 2012; Heo *et al*., 2012; Boseon Kim *et al*., 2015a; Gutiérrez-Vázquez *et al*., 2017). However, insignificant global changes in miRNA half-lives indicated that isomiR dynamics might not control miRNA expression levels as a general universal mechanism (Kingston and Bartel, 2019). In line with previous studies, control of miRNA turnover for specific miRNAs including the let-7 family members by TUT4/7-mediated 3′ uridylation is hard to refute and rule out (Heo *et al*., 2012; Thornton *et al.*, 2014; B. Kim *et al*., 2015). Moreover, NTA isomiRs such as the uridylated fraction have been shown to regulate non-canonical target repertoires and to increase the diversity of targets for a certain miRNA by mono- or di-nucleotide changes in its 3′ terminus (Yang *et al*., 2019). Furthermore, half-lives of miRNA are longer than their mRNA targets, with a median half-life of 25 hours in proliferating mouse embryonic fibroblasts (Kingston and Bartel, 2019). Therefore, NTA modifications of readily available stable miRNAs might be a means of fast response to the changing stimuli through targeting diverse repertoires of non-canonical mRNA targets. This probably provides a greater control on the overall physiological fate.

With this new knowledge of TUT4/7 biology, it is apparent that TUT4/7 are master regulators of the global transcriptome. Due to its involvement in the LIN28A:let-7 axis, special attention has been brought on the efficacy of targeting TUT4 and TUT7 to negatively impact cancer proliferation and progression (Piskounova *et al*., 2011; Lin and Gregory, 2015; Kim *et al*., 2020). Based on our findings, it is important to consider that inhibition of these uridylating enzymes comes together with compensatory and wide range of transcriptomic changes that vary depending on the cancer cell-type or status. Therefore, further understanding of and exploring such dynamics of gene regulation is crucial to avoid non-specific deleterious side-effects when targeting TUT4/7 for anti-cancer therapeutics.

## ACKNOWLEDGEMENTS

We thank members of the Gurdon Institute and Storm Therapeutics Ltd. for useful feedback and discussions. We thank Kay Harnish and the core NGS facility at the Gurdon Institute for help with processing RNA sequencing libraries. We are grateful to Sylviane Moss and Marc Ridyard for risk assessments and reagents orders, respectively. We are also grateful to Kin Man Suen, Rohan Sivapalan, Matylda Sczaniecka-Clift and Julia Coates for guidance on experimental protocols. We thank Jack Monahan, Alexandra Dallaire, Navin Brian Ramakrishna, Isabela Navarro and Archana Yerra for general discussion and feedback on paper drafts, and Dhiru Bansal for help with initial cell phenotyping assays. This work was primarily supported by STORM Therapeutics Limited, Cambridge, UK funding to R.M.; and by Cancer Research UK (C13474/A18583, C6946/A14492) and the Wellcome Trust (219475/Z/19/Z, 092096/Z/10/Z) grants to E.A.M. For Open Access, the author has applied a CC BY public copyright licence to any Author Accepted Manuscript version arising from this submission.

## AUTHOR CONTRIBUTIONS

Experimental design: R.M., A.S. and E.A.M.; Experimentation and method optimisation: R.M., G.F. and B.G.; Bioinformatic Analysis: J.P. and R.M.; Resources: R.M., A.S. and E.A.M.; Drafting the main text of the manuscript: R.M.; Figure design, visualization and preparation: J.P. and R.M.; Revision: R.M., J.P., G.F., A.S. and E.A.M. and Supervision: E.A.M..

## STAR Methods

### Key resource table

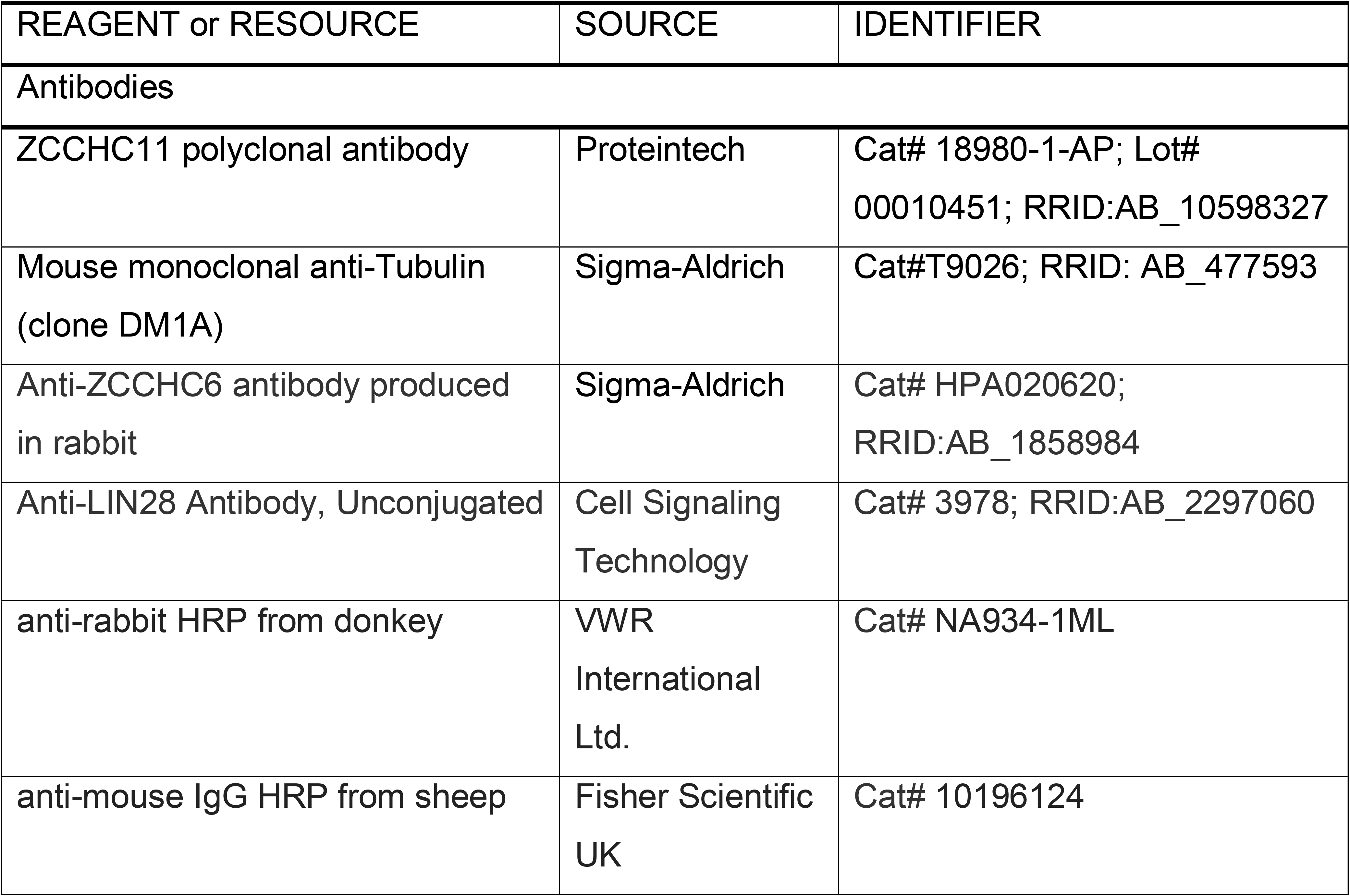
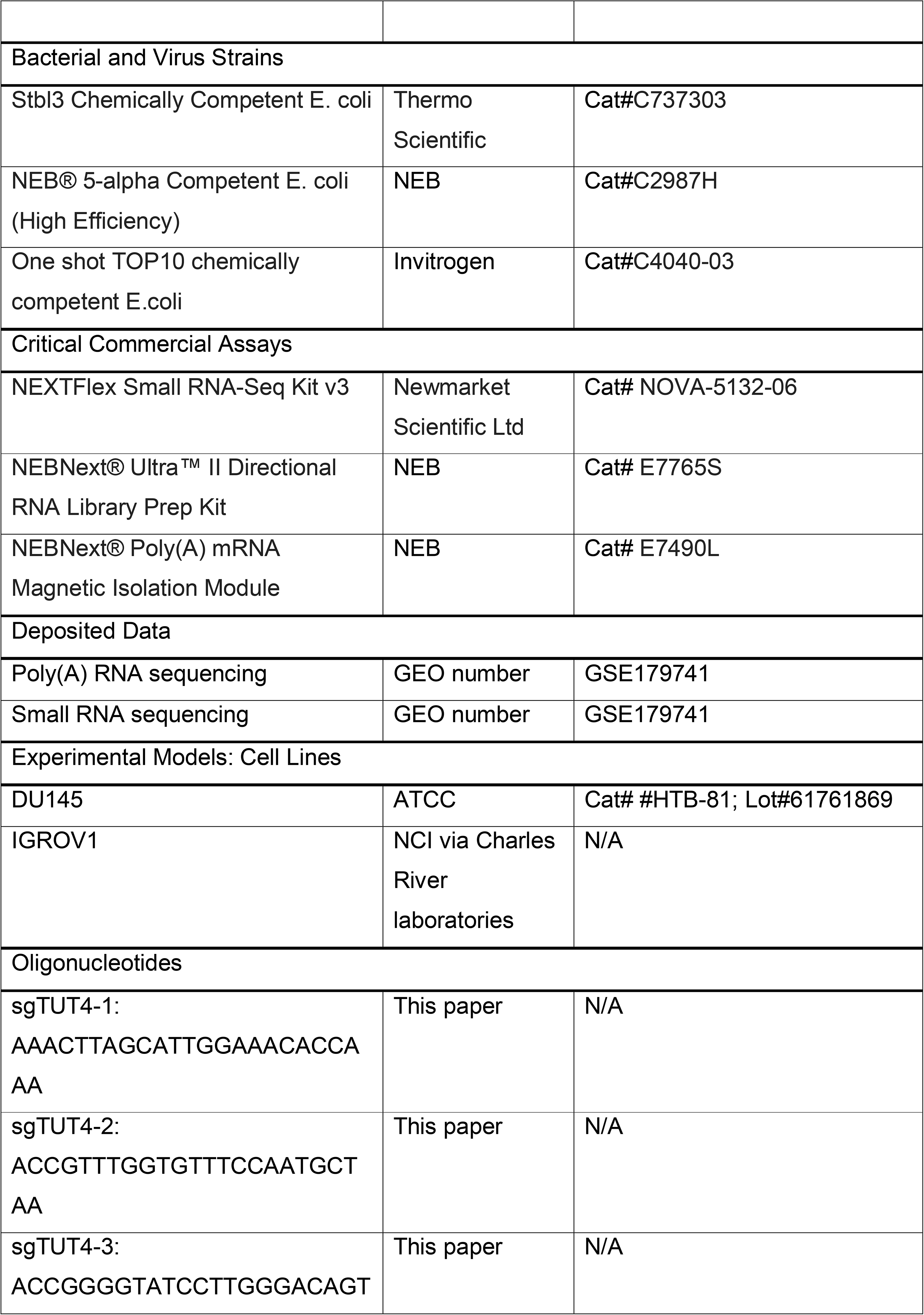
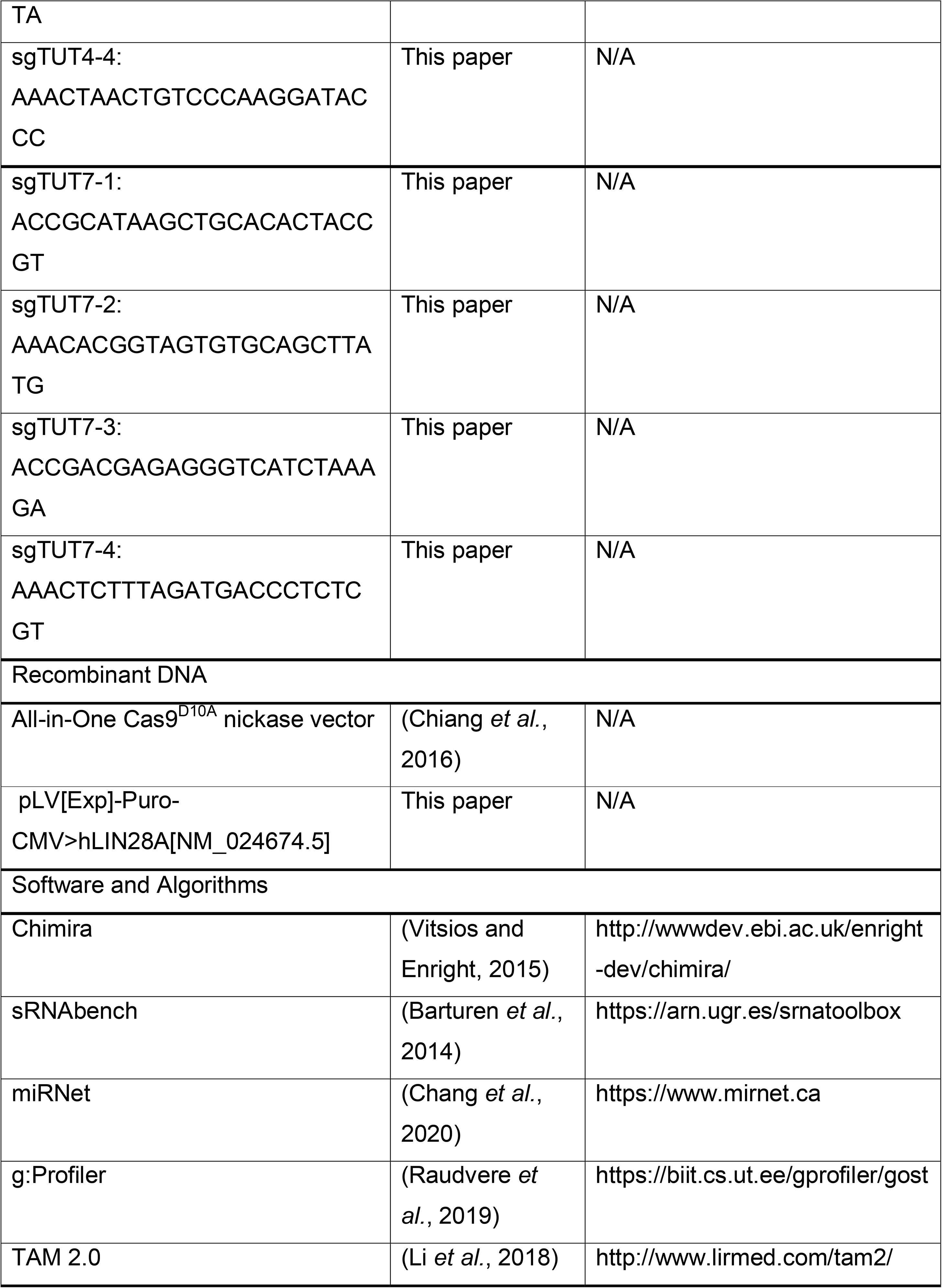

### Lead contact

Further information and requests for resources and reagents should be directed to and will be fulfilled by the Lead Contact, Eric A. Miska.

### Materials availability

Reagents generated for this study are available upon request.

### Data and code availability

The datasets generated during this study are available at NCBI’s Gene Expression Omnibus and are accessible through GEO Series accession number GSE179741.

### Method details

#### Cell culture

Prostate cancer cell line, DU145 (ATCC/LGC, Cat. No#HTB-81, Lot#61761869) and the ovarian cancer cell line IGROV1 (NCI via Charles River laboratories) were selected as the cell lines for use as *in vitro* models. DU145 complete media was made with the following reagents: EMEM (Gibco, Cat. No#41090093), 1X NEAA (Gibco, Cat. No#11140035), 10% FBS (Gibco, Cat. No#10270106, Lot#42G6287K), 1X Antibiotics (Life Technologies, Cat. No#15140122) and 5 ml Sodium Pyruvate (Life Technologies, Cat. No#11360-039). IGROV1 complete media comprised of the following: RPMI-1640 (Gibco, Cat. No#72400054), 1X NEAA (Gibco, Cat. No#11140035), 10% FBS (Gibco, Cat. No#10270106; Lot# 42G6287K), 1X Antibiotics (Life Technologies, Cat. No#15140122). Cell lines were maintained at 37°C, 5% CO2 in a humid incubator. Mycoplasma testing was performed using an EZ-PCR Mycoplasma Test Kit (Geneflow Ltd., Cat. No#K1-0210) according to the manufacturer’s recommendation. At 80-90% confluency, adherent cell cultures were dissociated using 1X Trypsin-EDTA (Life Technologies, Cat. No#25300054) and passaged at a 1:4 dilution. Icons to represent different tissues were created from BioRender.com.

#### Plasmid preparation and cloning

One shot TOP10 chemically competent cells (Invitrogen Ltd., Cat. No#C4040-03) were used and incubated at 37°C overnight post transformation according to the manufacturer’s recommendation. Following this, 4 ml of LB media was inoculated with a single colony and incubated at 37°C for 24 hours. Purification and extraction of plasmids for genotyping was performed using a PureLink Quick Plasmid Miniprep kit (Invitrogen, Cat. No#K210001). Plasmid sequences and their identity were validated by Sanger sequencing (Genewiz service) or using restriction enzymes. Endotoxin free plasmid preparation for transfection into cells was performed using a Qiagen Plasmid Maxi Kit (Cat. No#12163). Restriction enzymes required for cloning into the desired plasmid site were identified using CLC Workbench software. Successful digestions were confirmed using agarose gel electrophoresis. 1% agarose gel was prepared using 1 g of agarose powder (Bioline. Cat. No#BIO-41025) in 100 ml of 1X TBE buffer (Tris-borate-EDTA) and 1 μl of ethidium bromide solution (Merck, Cat. No#E1510). The digested products and 1KB Plus DNA ladder (Life Technologies, Cat. No#10787018) were loaded on separate wells of the agarose gel and run at a constant voltage of ∼130V for 40 minutes with 1X TBE as the buffer. Digested linear plasmid at the correct band size was cut and recovered from the gel using Zymoclean Gel DNA recovery kit (Zymo Research, Cat. No#D4002). Purity of the extracted DNA was checked using the 260 nm to 280 nm and the 260 nm to 230 nm wavelength absorbance ratios. If the ratio of absorbance at 260 nm to 230 nm was lower than 2.0, then a second round of purification was performed using a PureLink PCR purification kit (Life Technologies Ltd., Cat. No#K310002). Ligations at different vector/insert ratios were done overnight at 20°C using a T4 DNA ligase (NEB, Cat. No#M0202T).

#### Generation of catalytic knockouts

The protocol described in the following section is an adaptation of the method described in Chiang et al. (2016). Risk assessments and training prior to performing the gene editing experiments were completed and approved (Reference number EM68). TUT4 (ZCCHC11) Exon 12 and TUT7(ZCCHC6) Exon 14 were successfully targeted using CRISPR/Cas9-mediated gene editing. Cas9 expressing plasmid All-in-One nickase vector (Chiang et al., 2016) was used as the backbone vector (kind gift by Steve Jackson Laboratory, Gurdon Institute). This is an All-in-One vector that expresses a Cas9 mutant nickase (RuvC^D10A^), enhanced green fluorescent proteins (EGFP) and the target specific small guide RNAs. Two pairs of guide RNAs were ligated with T4 DNA ligase (NEB, Cat. No#M0202T) using restriction digestion using Bsa1 (NEB, Cat. No#R0535S) and BbsI (NEB, Cat. No#R0539S) digestion as per NEB recommendations. One shot TOP10 chemically competent cells (Invitrogen Ltd., Cat. No#C4040-03) were transformed and selected on LB plates with ampicillin selection. The transformed E. coli was grown in LB broth with ampicillin and the recombinant plasmids were purified using a Qiagen Plasmid Maxi Kit (Cat. No#12163). Plasmids were sent for Sanger sequencing (Genewiz service) to further validate the ligation and sequence of the sgRNA.

8 μg of endotoxin free plasmids were aliquoted in 30 μl of water. 10 cells were resuspended in 1 ml of resuspension buffer R (Neon Transfection System 100 μl kit-Thermo Scientific, Cat. No#MPK10025). 120 μl of cells in resuspension buffer were added to the plasmid mix. The DNA-cell mix was then electroporated in 3′ ml of electrolyte buffer E2 (Neon Transfection System 100 μ kit-Thermo Scientific, Cat. No#MPK10025) at 1400V for 30 milliseconds at the rate of 1 pulse. The cells were immediately transferred to a serum rich media. EGFP expression after 48 hours of electroporation denoted successful transfection.

Cells were trypsinized and washed with phosphate buffered saline (PBS). Centrifugation was done at 1000 rpm for 3 minutes to pellet the cells. The cell pellet was resuspended in 500 μl of PBS. EGFP-positive cells were either sorted in a single-cell suspension per well of a 96-well plate or bulk sorted in a single tube with serum-rich media. Bulk sorted control cells and EGFP positive cells were plated in a 10 cm petri-dish with serum-rich media. Crude cell lysates were prepared using 5 μl of cell suspension in 20 μl of DirectPCR lysis reagent containing Proteinase K (Viagen Biotech, Cat. No#302-C) by following manufacturer instructions. Primers were designed either ∼300 bp or ∼600 bp away from the site targeted by the sgRNAs. PCR was performed using Q5 high fidelity polymerase (NEB, Cat. No#M0491) using recommended settings. Clones with unexpected band shifts as compared to the wildtype were then sent for Sanger sequencing to confirm deletions and insertions. Genomic DNA for genotyping was purified using a DNeasy Blood & Tissue kit (Qiagen, Cat. No#69504). Mutations on each copy of the gene were further explored using a TOPO TA Cloning kit (Invitrogen, Cat. No#450030) according to the manufacturer’s recommendations. Further validation was done via western blotting

#### Stable cell line generation

The lenti-X 293T cell line was kindly provided by the Steve Jackson lab (Gurdon Institute). A risk assessment (Reference Number EM67) was completed and approved before performing experiments that used lentiviruses. psPAX2 (Addgene Plasmid #12260) that expresses gag and pol genes along with pMD2.G (Addgene Plasmid #12259) that expresses the VSV-G envelope protein were used as packaging plasmids. Validation of psPAX2 was done with EcoRI (NEB, Cat. No#R3101), which results in two bands at ∼6 kb and ∼4 kb on agarose gel electrophoresis. Validation of pMD2.G was done by double digestion with BamH1 (NEB, Cat. No#R3136) and HindIII (NEB, Cat. No#R3104) which results in two bands at ∼3 kb and ∼2.8 kb on agarose gel electrophoresis. Transfer vectors used for viral production are provided in Table 2.2. All transfer vectors were driven by the CMV promoter and had puromycin as the antibiotic selective marker.

The protocol for lentiviral production was obtained from the Steve Jackson Laboratory (Gurdon Institute). For virus production, 10 cm petridishes (Corning, Cat. No#10212951) were pre-coated with 0.1% gelatin for 15 minutes. The lenti-X 293T cells were seeded at 1:3 and 1:4 dilutions on two gelatin coated plates. The following day, the cell cultures at 80-90% confluency were chosen for lentiviral production. 7.23 μg of psPAX2, 1.56 μg of pMD2.G and 2.93 μg of the transfer plasmid with the transgene of interest were aliquoted in a 1.5 ml Eppendorf tube and thoroughly mixed by pipetting (Tube 1). 624 μl of Opti-MEM (Gibco, Cat. No#11058021) and 35.1 μl of TransIT-Lenti transfection reagent (Mirus Bio, Cat. No#MIR6600) were aliquoted in another 1.5 ml Eppendorf tube and thoroughly mixed (Tube 2). Both Tube 1 and Tube 2 were incubated for 5 minutes at room temperature in a biosafety cabinet in a Containment Level 2 room. Subsequently, the contents of Tube 2 were added and mixed with Tube 1 and incubated for 25-30 minutes at room temperature. After the incubation, the lenti-X 293T cells were transfected with the mix and lentiviruses from the condition media were collected after 48 hours. The collected viruses were syringed through a 0.45 μm filter (Sartorius, Cat. No#16555-K). The harvested lentiviruses were then stored at 4°C for up to a week or at -70°C for longer storage.

Recipient cells were passaged at 1:2, 1:3 and 1:4 dilution. Cell cultures at 60-70% confluency were chosen for transduction. Cells were initially transduced with a range of viral doses in 5-fold increments starting at 10 μl. The maximum viral dose was ∼500 μl per well. Complete media was used as the dilution media. Transduced cells were incubated for 48 hours.

Puromycin (InvivoGen, Cat. No#ant-pr-1) was used as the antibiotic selective marker. Prior to transduction, the optimal puromycin dose was determined by incubating cells at 0, 0.5, 1, 1.5, 2 and 4 μg/ml of puromycin diluted in complete media. Cells were incubated for a period of 10 days with media replacement every 3-4 days. The minimum puromycin concentration that was lethal for 100% of the non-infected cells was selected as the optimal puromycin selection concentration for that cell line. After transduction, infected cells were trypsinised and seeded in a new well of a 6-well plate along with a non-infected control in a separate well and incubated with the optimal puromycin dose for 10 days with media replacement every 3-4 days. The non-transduced control did not survive after 10 days but the cells with a successful integration event survived the puromycin selection and treatment. Stable cell lines were further validated for the presence of the transgene on a western blot or under a fluorescence microscope if the transfer plasmid had an EGFP or mCherry marker. Varying degrees of success on generating stable cell lines were observed depending on the transgene size.

#### Western blot

Precast NuPAGE 3-8% Tris-acetate gels (Invitrogen, Cat. No#EA0378BOX) and NuPAGE 4-12% Bis-Tris Gel (Invitrogen, Cat. No#NP0335BOX) were used for separating the proteins as per manufacturer instructions. Running buffers, NuPAGE TA SDS (Invitrogen, Cat. No#LA0041), NuPAGE MOPS SDS (Invitrogen, Cat. No#NP0001) and NuPAGE Transfer buffer (Invitrogen, Cat. No#NP00061) were used. Primary antibodies used were ZCCHC11 (Proteintech, Cat. No#18980-1-AP, Lot#00010451), ZCCHC6 (Proteintech, Cat. No#25196-1-AP), ZCCHC6 (Merck, Cat. No#HPA020620), LIN28A (Cell Signalling, Cat. No#3978), LIN28B (Cell Signaling, Cat. No#4196) and Tubulin (Merck, Cat. No#9026). Secondary antibodies used were anti-rabbit (GE Healthcare, Cat. No#NA934) and anti-mouse (GE Healthcare, Cat. No#NA931).

#### RNA extraction and quality control

RNA was isolated using TRISure (Bioline, Cat. No#BIO-38033) and phase lock tubes (Quantabio, Cat. No#733-2478) following standard chloroform-isopropanol extraction. Isolated RNA then underwent TURBO DNase treatment (Ambion, Cat. No#AM1907) to remove DNA contaminants. Purity values were measured using Nanodrop. Qubit RNA BR Assay kit (Thermofisher Scientific, Cat. No#Q10210) was used to quantify RNA. Reverse transcription was performed using appropriate TaqMan kits (Applied Biosystems) and qPCR was done as per instructions. Analysis of the qPCR results and statistics were performed using the R package ‘pcr’ (Ahmed and Kim, 2018).

#### Small RNA library preparation and analysis

The small RNA libraries were prepared using a NEXTFlex Small RNA-Seq Kit v3 3′ (Cat. No#NOVA-5132-06) according to the manufacturer’s recommendations. The adapters, TGGAATTCTCGGGTGCCAAGG were trimmed using cutadapt (Martin, 2011) and further processed to trim the first and last four random bases that act as unique molecular identifiers or UMIs. Use of such UMIs ensures the identification of PCR duplicates th a t arise as a result of PCR amplification steps. Quality checks were performed using FastQC (Andrews, 2010) to determine if miRNAs had a clear peak at the expected size. Small RNA reads were analysed with miRDeep2 (default parameters; Friedla□nder et al., 2012) using the *H. sapiens* (hg38) reference genome. Counts were imported into R and differential expression analysis was performed with DESeq2 (FDR <0.01; Love et al., 2014). Principal component analysis (PCA) in R was performed to determine if samples of the same genotype clustered together. K-means clustering (k = 15) of miRNAs was performed to group miRNAs with similar expression patterns across the samples. miRNA set analysis was performed with TAM 2.0 (Li et al., 2018). For proportion of modifications such as SNPs, analysis was performed using Chimira software (Vitsios and Enright, 2015) and counts were obtained. All subsequent analyses of the modification profiles and counts sRNAbench (Barturen et al., 2014) software. Ratios or percentages of non-templated additions were calculated by dividing the modified reads by the total number of reads (unless specified otherwise). Statistical analysis was performed using the ggstatsplot package in R (Patil, 2018).

#### mRNA library preparation and analysis

The mRNA sequencing libraries were prepared using NEBNext® UltraTM II Directional RNA Library Prep kits (NEB, Cat. No#E7765S) with the oligo dT selection module ac-cording to the manufacturer’s recommendation. Firstly, raw reads were trimmed for adaptors, low quality sequences and short reads with Trimmomatic (parameters: ILLUMINACLIP: TruSeq3-SE.fa:2:30:10 SLIDING-WINDOW:4:28 MINLEN:20; Bolger et al., 2014). For the second step, ribosomal RNAs were removed with sortmeRNA (default parameters; Kopylova et al., 2012). Following this, read mapping was done to the reference genome *H. sapiens* (hg38) with HISAT2(default parameters; Kim et al., 2015b) and reads were, then, counted on genes with HTSeq-count (Anders et al., 2015). Counts were imported into R and differential gene expression analysis was performed with DESeq2 (FDR <0.01; Love et al., 2014). Gene Ontology analysis of differentially expressed genes was performed with g:Profiler (Raudvere et al., 2019). Further, statistical analyses on normalised counts for the boxplots were performed using the ggstatsplot package in R (Patil, 2018). The web-based platform miRNet 2.0 (Chang *et al*., 2020) was utilized to identify miRNA-mRNA interactions. The list of IGROV1-specific downregulated mRNAs and upregulated miRNAs as query or vice versa and gene ontology terms related to wound healing or cell migration were selected under function explorer (Algorithm-Hypergeometric test; Database-GO:BP).

#### Cell proliferation assays

Several dilutions of the DU145 wildtype and mutant cell lines (200, 400, 600, 800, 1000 and 1200 cells/well) were seeded in a 6-well plate. Plates were incubated at 37°C and 5% CO2 in a humid incubator for 15 days. For IGROV1, 50,000/10,000 cells/well in a 24-well plate were seeded and incubated for 4 days. At the endpoint, plates were washed and stained with Crystal Violet (Merck, Cat. No#V5265-250ML). In addition, proliferation assays at different dilutions and plate formats were performed using a Sartorius/Essen Biosciences IncuCyte^®^ System. For the 96-well plate format, the cell dilutions for each well per genotype were 10,000, 20,000 or 30,000 in technical replicates of three. Confluency was determined by label-free time lapse imaging at 2-hour intervals using the IncuCyte^®^ System.

#### Growth curve measurements

Raw proliferation data (Percent confluency) was obtained from time-lapse imaging using the IncuCyte^®^ analysis software. Data were analysed using the growthcurver R package (Sprouffske and Wagner, 2016). Initial confluency was subtracted from the confluency value at each timepoint. The growth rate was obtained by fitting the data to a logistic equation as given below (Sprouffske and Wagner, 2016).

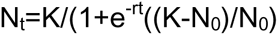

In this logistic equation, *N*0 and *Nt* are cell confluency at the beginning and at time *t*, respectively. *K* is the maximum carrying capacity and *r* is the growth rate.

#### Scratch assay

Cells were seeded in IncuCyte^®^ ImageLock 96-well plates (Sartorius, Cat. No# 4806). At 90-100% confluency, a uniform scratch was introduced using the IncuCyte^®^ Cell Migration Kit (Sartorius, Cat. No#4493) and following manufacturer’s instructions. Wound closure was measured by time lapse imaging at 2-hour interval and results were analysed using the IncuCyte^®^ system. The R code for analysing data was obtained from Grada et al. (2017).

#### Trans-well migration assay

8.0 μ m pore 6.5 mm Corning Trans-well inserts in a 24-well format (Merck, Cat. No#CLS3422-48EA) were used for a qualitative cell migration assay. For the experimental group, the lower chamber comprised of 800 μl of cell culture medium supplemented with 10% fetal bovine serum (FBS) (Life Technologies, Cat. No#10270106, Lot#42G6287K). For the control group, cell culture media without FBS was pipetted into the lower chamber. Adherent cells were trypsinised with 1X Trypsin (Life Technologies, Cat. No# 25300054) and harvested in media without FBS. 150 μ of cell dilutions with 5 × 10 cells/ml for DU145 were pipetted carefully into the upper insert chamber. For IGROV1, 100,000-150,000 cells were pipetted into the upper insert chamber. Trans-well plates were incubated for 24 hours at 37°C and 5% CO2 in a humid incubator. At the end point, plates were washed twice in 1X PBS and stained with Crystal Violet (Merck, Cat. No#V5265-250ML) for 10 minutes. After staining, inserts were washed with water. The non-migrated cells on the inside of the insert were removed with a cotton swab. In summary, this protocol was adapted from an application note entitled ‘Cell migration and invasion quantification assay with acetic acid-dependent elution of crystal violet’ by Corning Incorporated, Life Sciences, Shanghai, China.

## SUPPLEMENTARY LEGENDS

**Figure S1: Related to Figure 1.**
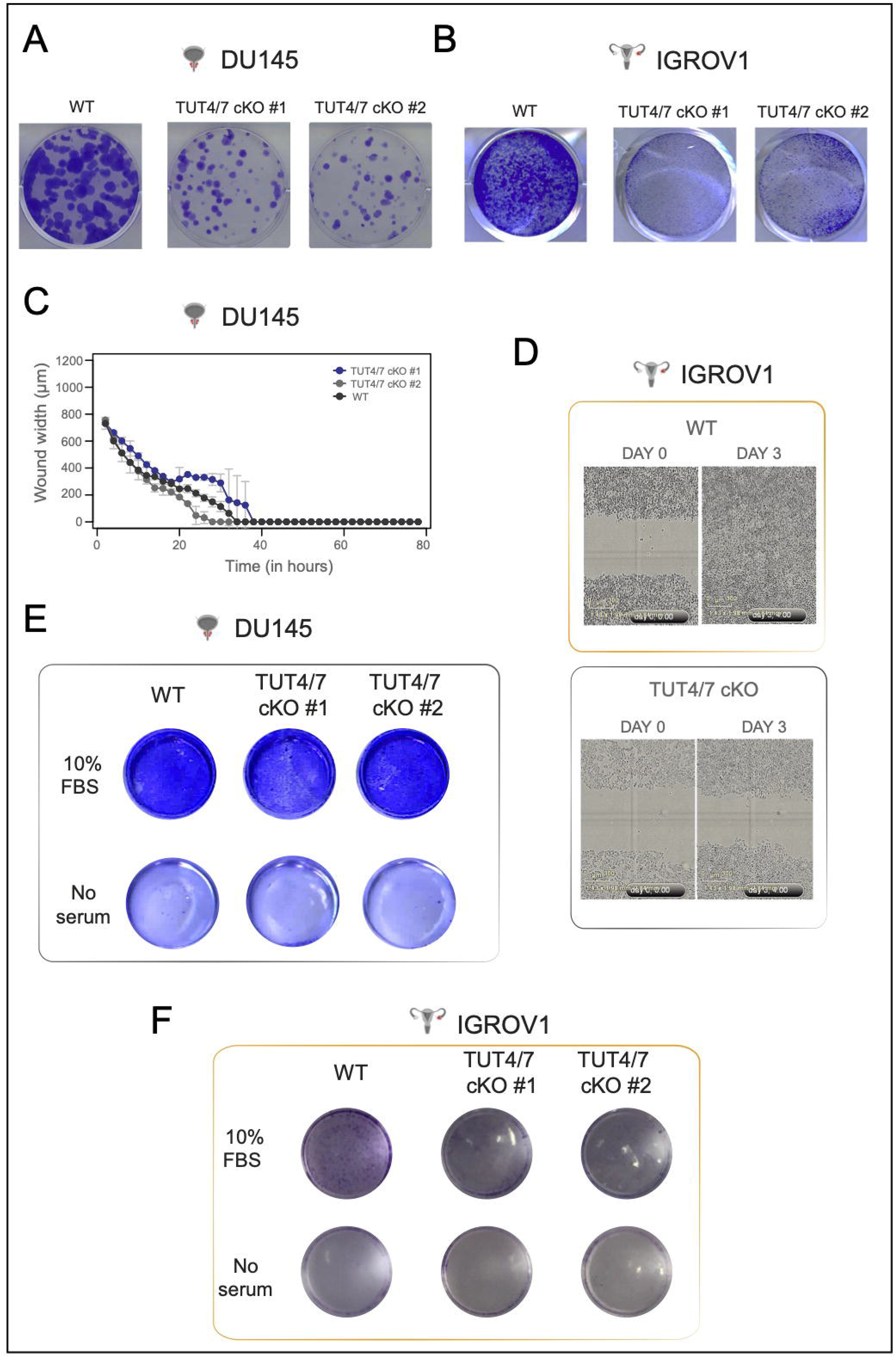
Cell proliferation and migration properties of TUT4/7 double mutants. (A) Crystal violet staining at the end of a colony formation assay. Number of cells at the start = 400. End day number = 15; number of replicates, n=3; TUT4/7 cKO #1 is one independent TUT4/7 double mutant clone and TUT4/7 cKO #2 is the other. WT denotes the wildtype control. (B) Crystal violet staining at the end of cell proliferation assay. End day number = 4; number of replicates, n = 2; TUT4/7 cKO #1 is one independent TUT4/7 knockout clone and TUT4/7 cKO #2 is the other. WT denotes the wildtype control. (C) Quantitative wound healing assays with measurements at 2-hour intervals using the Incucyte^®^ systems. The y-axis denotes the wound width in μm, and the x-axis represents time in hours (n=2). Cell line = DU145. WT is the control DU145 cell line. TUT4/7 cKO #1 and TUT4/7 cKO #2 are the TUT4/7 double mutants. (D) Brightfield images of TUT4/7 double mutants (TUT4/7 cKO) of IGROV1 and the wildtype control (WT) on day 0 (start point) and day 3′ (end point) of the scratch assay. (E and F) Trans-well cell migration assay for DU145 TUT4/7 double mutants (E) and IGROV1 TUT4/7 double mutants (F) with WT as their respective controls. TUT4/7 cKO #1 and TUT4/7 cKO #2 are the TUT4/7 double mutants (n=2).

**Figure S2: Related to Figure 2.**
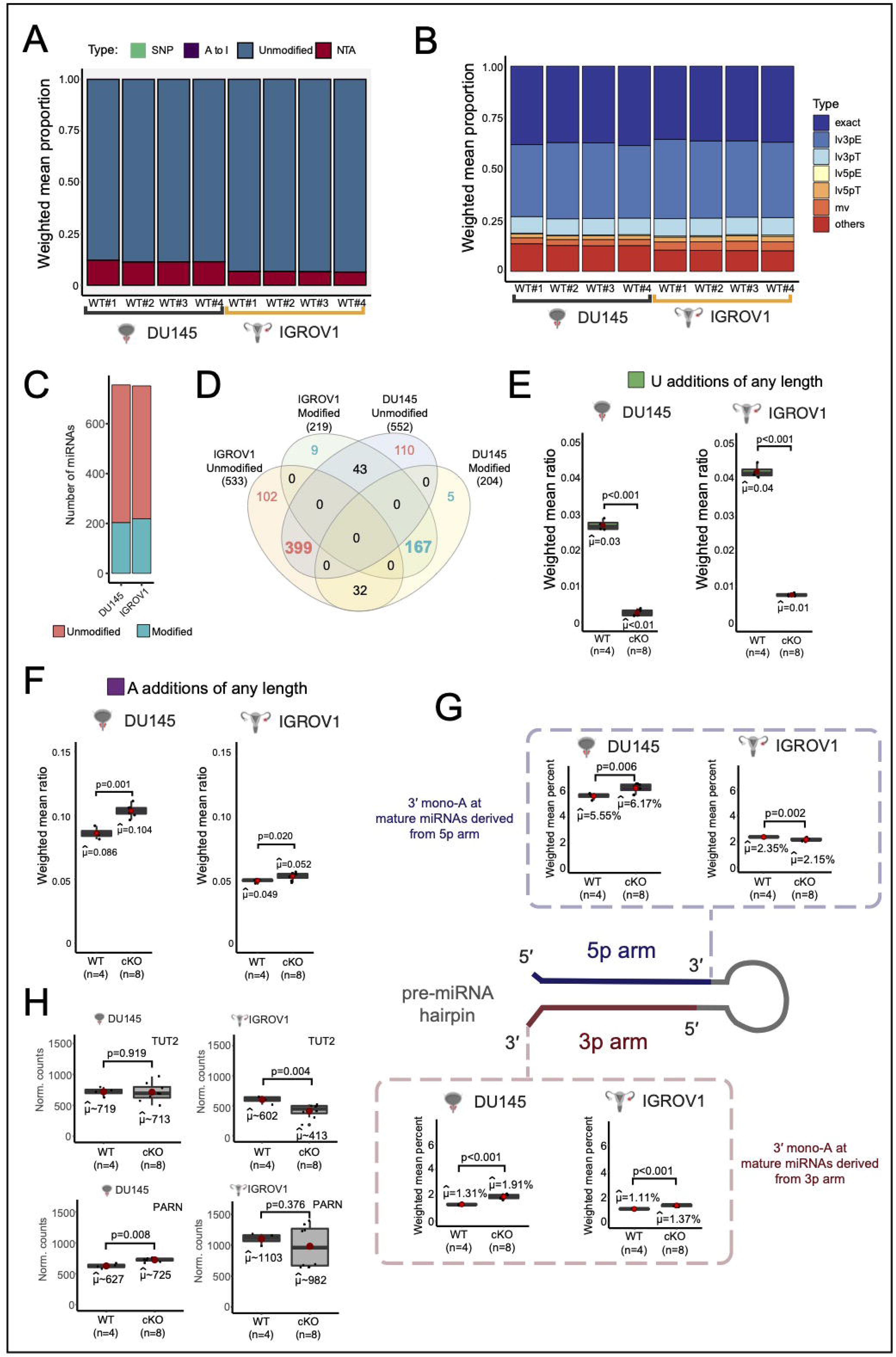
Proportion of modified isomiRs in DU145-derived and IGROV1-derived cell lines. (A) WT#1, WT#2, WT#3 and WT#4 are the four biological replicates of either DU145 denoted by the grey line or of IGROV1 denoted by the yellow line. Weighted mean proportion is the ratio of unmodified or modified reads specific to the type of modification (A to I, SNP or NTA) to the total number of reads mapping to miRNAs in the sample. (B) The miRNA population is dominated by miRNAs with variations in their 3′ end. Canonical miRNAs which exactly match their template are represented by “exact”; miRNAs with 3 templated extensions are denoted by “lv3pE”; miRNAs with a trimmed 3′ end are indicated by “lv3pT”. Similarly, miRNAs with 5 templated extensions are denoted by “lv5pE” and miRNAs with a trimmed 5′ end are indicated by “lv5pT”. “mv” denotes multi-length variants which do not match both the 5′ end and 3′ end. The rest of the miRNAs are placed in “others”. (C) Bar chart with the number of modified and unmodified 3p mature miRNAs for DU145 and IGROV1 on the y-axis. (D) Venn diagram showing overlaps of modified and unmodified 3p mature miRNAs between the two cell lines. (E) Non-templated U-additions of any length in the wildtype control (WT; n=4) and TUT4/7 double mutants (cKO; n=8) of DU145 and IGROV1. μ cap denotes the mean and the statistics used are described at the bottom of each boxplot. (F) Non-templated A-additions of any length in the wildtype control (WT; n=4) and TUT4/7 double mutants (cKO; n=8) of DU145 and IGROV1. μ cap denotes the mean and the statistics used are described at the bottom of each boxplot. (G) Mono-A additions at 3p and 5p mature miRNAs in the wildtype control (WT; n=4) and TUT4/7 double mutants (cKO; n=8) of DU145 and IGROV1. μ cap denotes the mean and p-value is indicated by p. (H) The y-axis indicates normalised counts. The expression counts depict changes in the exonucleases PARN along with the adenylating TENT, TUT2 in the prostate cancer cell line DU145. WT = Wildtype control (n=4) and cKO = TUT4/7 double mutants (n=8). p-value (p) is denoted at the bottom of each plot. μ cap indicates the mean.

**Figure S3: Related to Figure 3.**
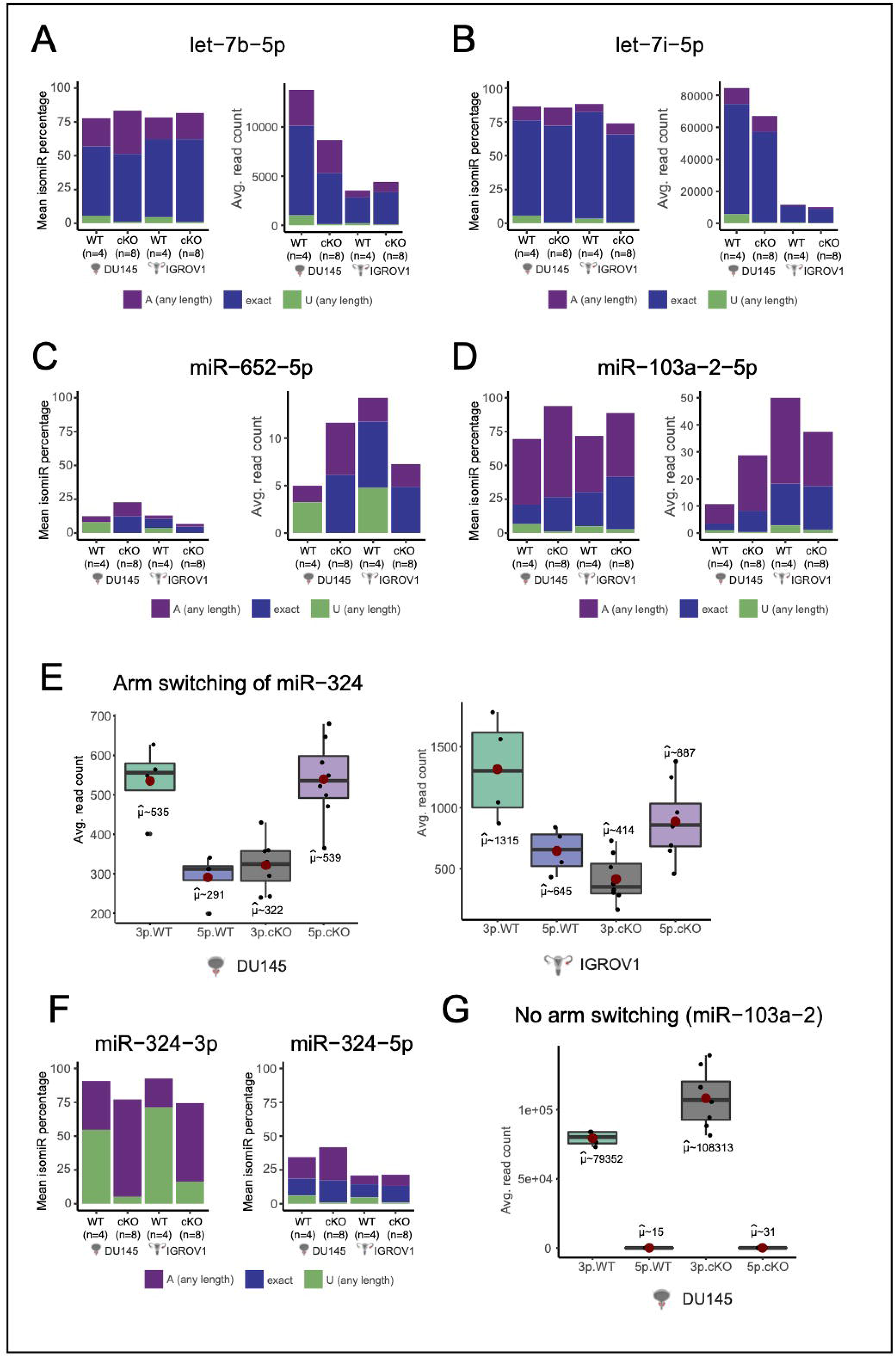
TUT4/7-mediated variations in isomiR populations in select miRNAs. (A) Proportion and abundances of canonical, adenylated and uridylated isomiRs of let-7b-5p. (B) Proportion and abundances of canonical, adenylated and uridylated isomiRs of let-7i-5p. (C) Proportion and abundances of canonical, adenylated and uridylated isomiRs of miR-652-5p. (D) Proportion and abundances of canonical, adenylated and uridylated isomiRs of miR-103a-2-5p. (E) 5p/3p arm switching of miR-324 in the WT and TUT4/7 double mutants (cKOs) in DU145 and IGROV1. (F) Proportion of canonical, adenylated and uridylated isomiRs of miR-324-3p and miR-324-5p. (G) Abundances of 5p and 3p mature miRNAs of miR-103a-2 in the prostate cancer cell line DU145 (WT) and its corresponding TUT4/7 double mutants (cKOs).

**Figure S4: Related to Figure 4.**
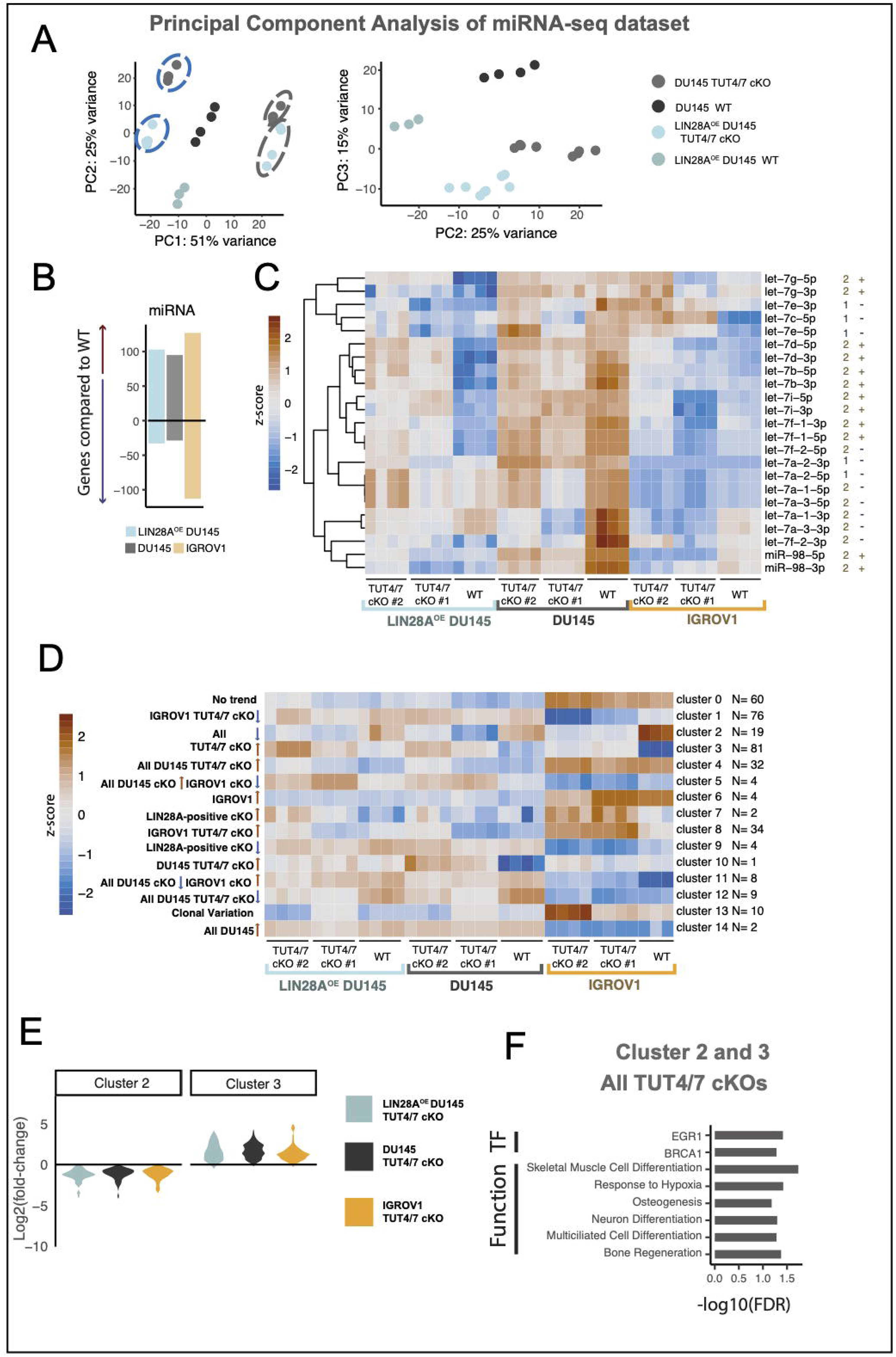
Analysis of the miRNA deregulation in TUT4/7 double mutants relative to the control. (A) Principal component analysis of miRNA dataset. PC1 versus PC2 and PC2 versus PC3 in miRNA data for DU145 prostate cancer cell line and related genotypes. The two different clones of the same genotype are highlighted by blue and grey circles. (B) Global miRNA deregulation trend with LIN28A*^OE^* DU145 in blue, DU145 in grey and IGROV1 in yellow. (C) Heatmap representing expression levels of the let-7 family members in the TUT4/7 double mutant clones (TUT4/7 cKO #1 and TUT4/7 cKO #2) and the wildtype controls (WT) of LIN28A*^OE^* DU145, DU145 and IGROV1. Each column represents a biological replicate. The let-7 miRNAs belonging to Group I are denoted by “1” and those belonging to Group II are denoted by “2”. “+” indicates that the precursor of the let-7 miRNA contains the (U)GAU sequence motif for LIN28A binding via its CSD domain and “-” indicates the absence of this binding motif. The colour key on the left indicates the z-score or scaled normalised expression from high (in brown) to low (in blue). (D) K-means (k=15) clustering of all miRNAs except the let-7 family groups the miRNAs into 14 distinct groups with similar expression patterns. The TUT4/7 double mutant clones are represented by TUT4/7 cKO #1 and TUT4/7 cKO #2 while the wildtype control of LIN28A*^OE^* DU145, DU145 and IGROV1 is denoted by WT. Each column represents a biological replicate. The colour key on the left indicates the z-score or scaled normalised expression from high (in brown) to low (in blue). (E) Violin plots displaying the expression trends in Clusters 2 and 3. Fold change is calculated in the TUT4/7 double mutants (TUT4/7 cKO) relative to their respective wildtype control. (F) miRNA set enrichment analysis was performed with TAM 2.0 (Li et al., 2018). Enriched associations for the miRNA Cluster 2 and Cluster 3′ with a similar deregulation pattern in TUT4/7 double mutants independent of the cell line of origin.

**Figure S5: Related to Figure 5.**
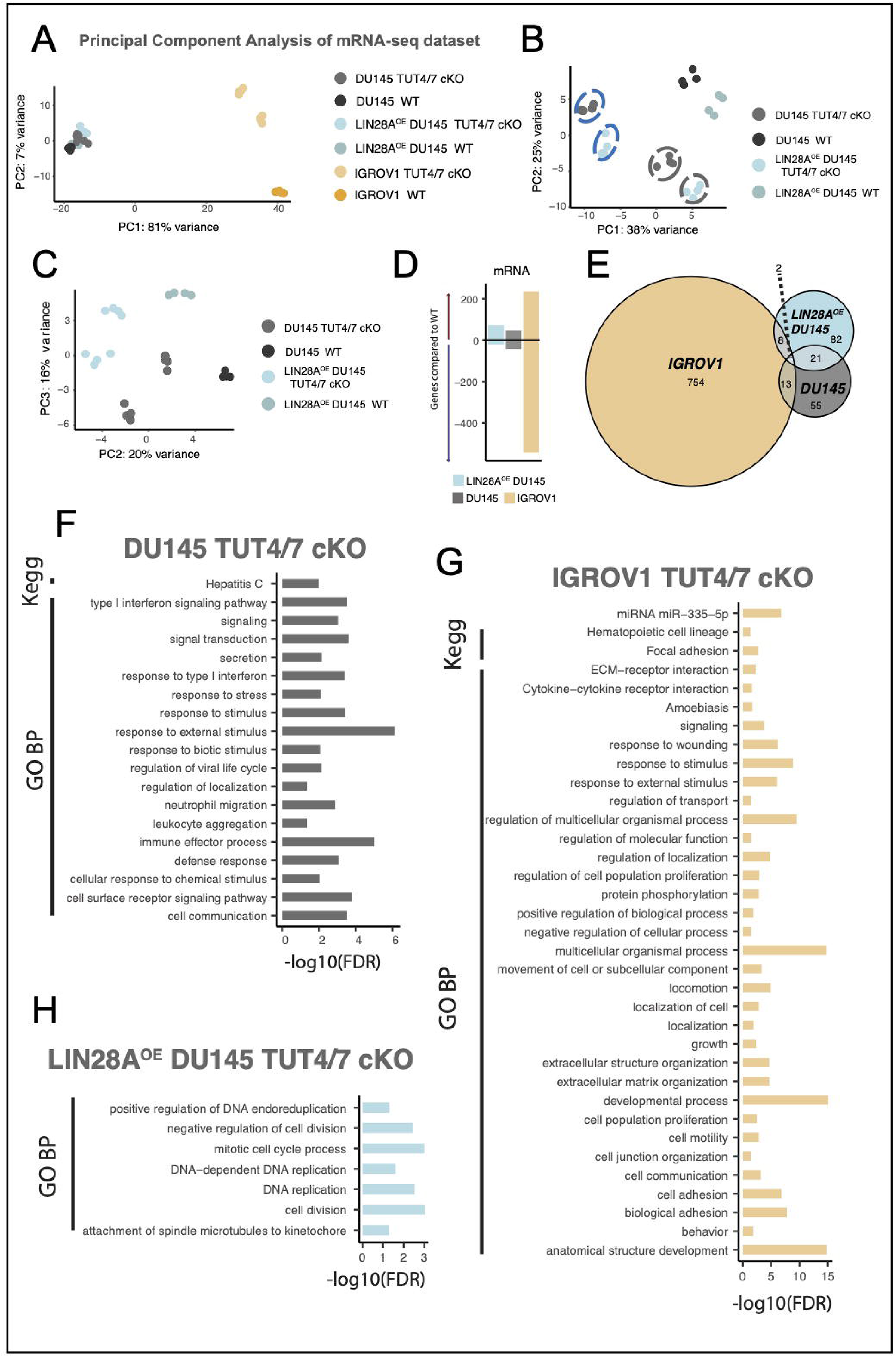
Deregulated mRNAs in TUT4/7 double mutants. (A, B and C) Principal component analysis of mRNA dataset. PC1 versus PC2 in mRNA data for all genotypes (A), PC1 versus PC2 in mRNA data for DU145 prostate cancer cell line and related genotypes (B), and PC2 versus PC3 in mRNA data for the DU145 prostate cancer cell line and related genotypes (C). The two different clones of the same genotype are highlighted by blue and grey circles. (D) Global mRNA deregulation trend with LIN28A*^OE^* DU145 in blue, DU145 in grey and IGROV1 in yellow. (E) Venn diagram showing gene overlap. (F, G and H) Gene Ontology analysis of differentially expressed genes was performed with g:Profiler (Raudvere et al., 2019). Enriched terms for the differentially regulated mRNAs in DU145 TUT4/7 double mutants (F). Enriched terms for the differentially regulated mRNAs in IGROV1 TUT4/7 double mutants (G). Enriched terms for the differentially regulated mRNAs in LIN28A*^OE^* DU145 TUT4/7 double mutants (H).

**Figure S6: Related to Figure 5.**
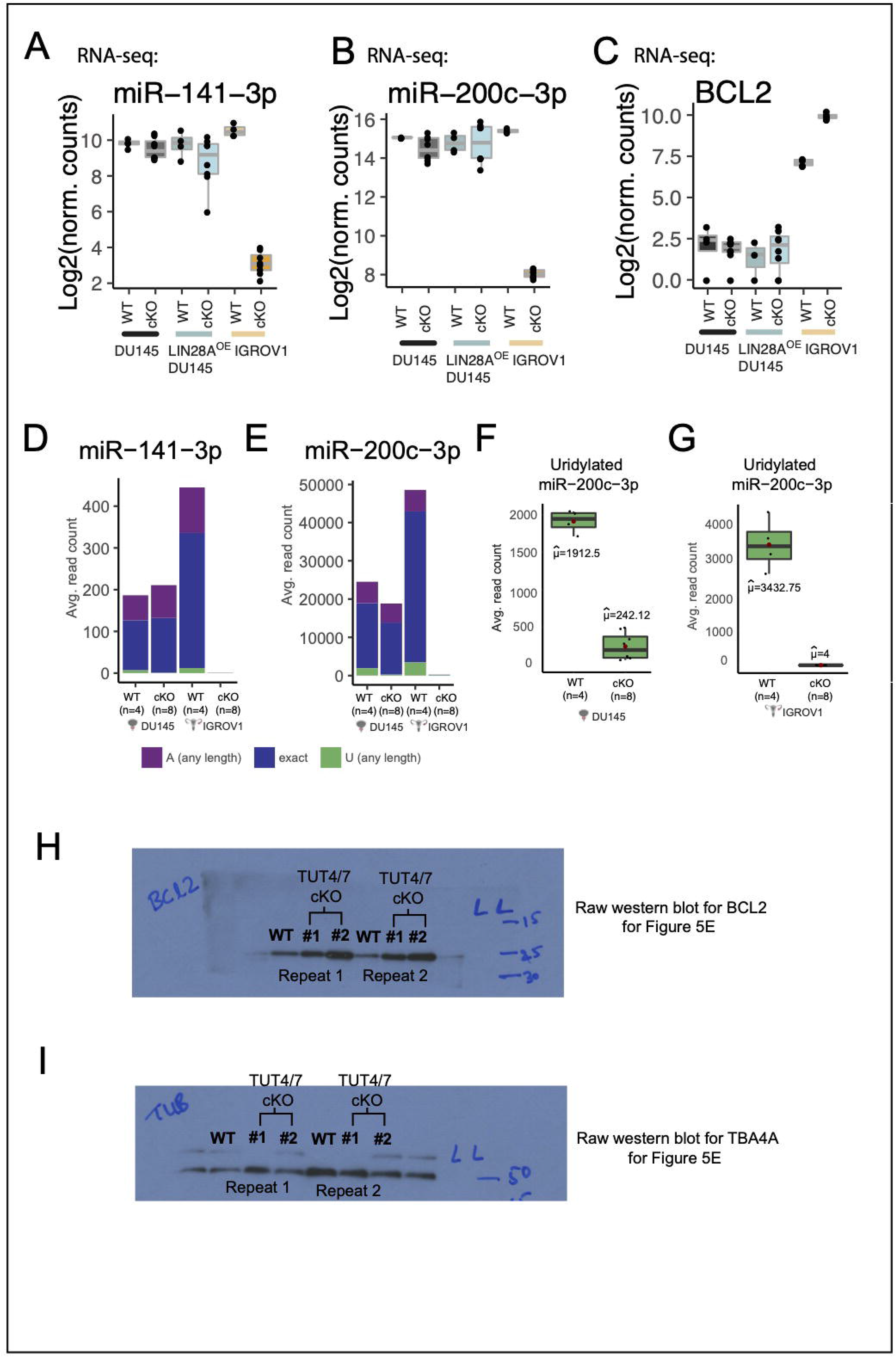
miR-200c cluster and BCL2 in TUT4/7 double mutants. (A, B) Expression levels of miR-141-3p (A) and miR-200c-3p (B) in DU145, LIN28A*^OE^* - DU145, IGROV1 and derived TUT4/7 cKOs from RNA seq data analysed with miRDeep2 (Friedla□nder et al., 2012). Number of biological replicates (n) for WT = 4. TUT4/7 cKO #1 represents one independent clone (n=4) and TUT4/7 cKO#2 (n=4) represents the other. (C) Expression levels of BCL2 in DU145, LIN28A*^OE^* DU145, IGROV1 and derived TUT4/7 cKOs from RNA-seq data. Number of biological replicates (n) for WT = 4. TUT4/7 cKO #1 represents one independent clone (n=4) and TUT4/7 cKO #2 (n=4) represents the other. (D, E) Abundances of canonical, adenylated and uridylated isomiRs of miR-141-3p (D) and miR-200c-3p (E) in DU145, IGROV1 and derived TUT4/7 cKOs from RNA-seq data analysed with sRNAbench (Barturen et al., 2014). (F) Abundances of uridylated isomiRs of miR-200c-3p in DU145 and derived TUT4/7 cKOs from RNA-seq data analysed with sRNAbench software (Barturen et al., 2014). (G) Abundances of uridylated isomiRs of miR-200c-3p in IGROV1 and derived TUT4/7 cKOs from RNA-seq data analysed with sRNAbench software (Barturen et al., 2014). (H) Raw western blot image for BCL2 for Figure 5E. (I) Raw western blot image for TBA4A for Figure 5E.

---

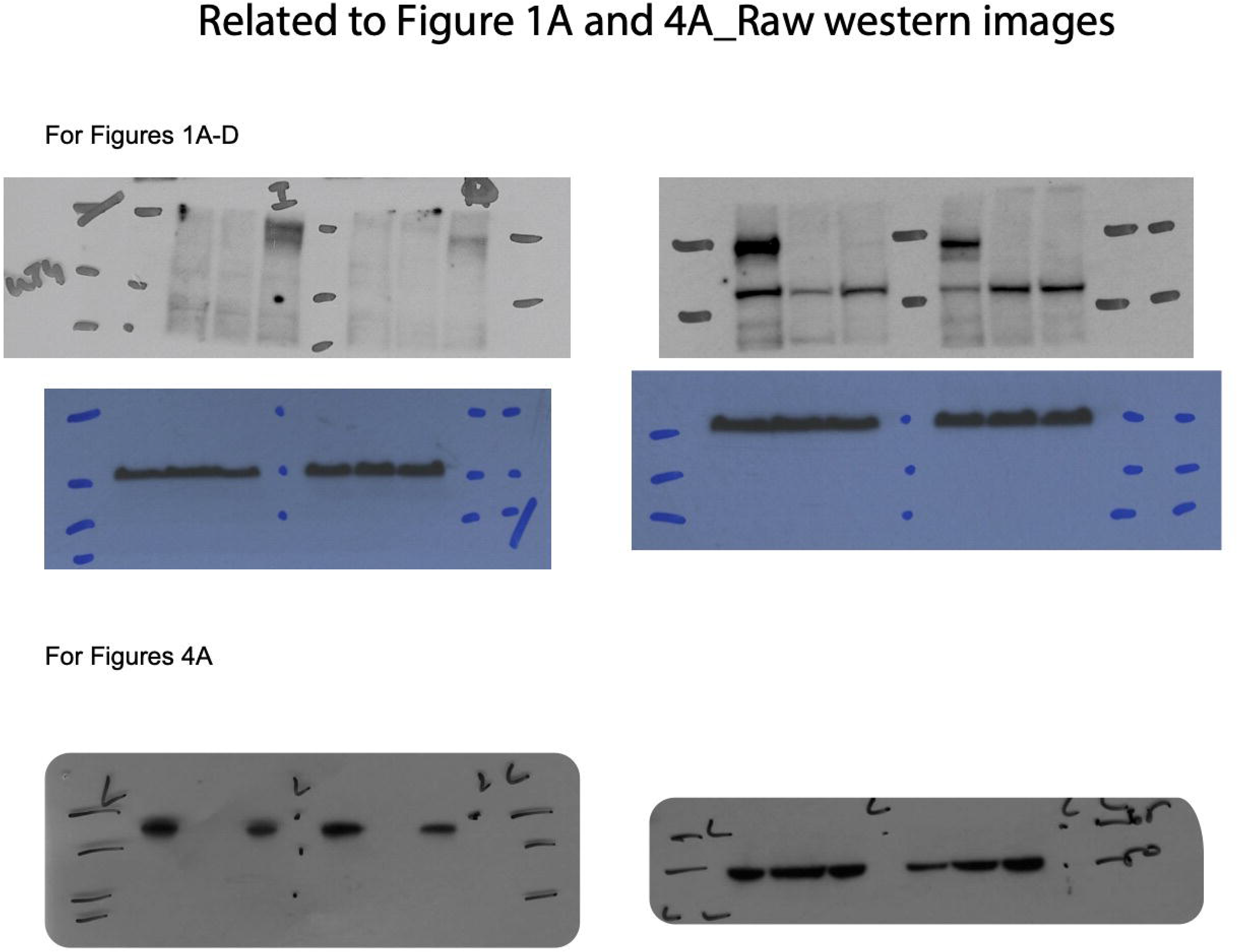

## BIBLIOGRAPHY

Ahmed, M., & Kim, D. R. (2018). pcr: an R package for quality assessment, analysis and testing of qPCR data. PeerJ, 6, e4473.

Bartel, D. P. (2004). MicroRNAs: genomics, biogenesis, mechanism, and function. cell, 116(2), 281–297.

Bartel, D. P. (2018). Metazoan micrornas. Cell, 173(1), 20–51.

Barturen, G., Rueda, A., Hamberg, M., Alganza, A., Lebron, R., Kotsyfakis, M., … & Hackenberg, M. (2014). sRNAbench: profiling of small RNAs and its sequence variants in single or multi-species high-throughput experiments. Methods in Next Generation Sequencing, 1(1), 21–31.

Bénard, J., Da Silva, J., De Blois, M. C., Boyer, P., Duvillard, P., Chiric, E., & Riou, G. (1985). Characterization of a human ovarian adenocarcinoma line, IGROV1, in tissue culture and in nude mice. Cancer research, 45(10), 4970–4979.

Burroughs, A. M., Ando, Y., de Hoon, M. J., Tomaru, Y., Nishibu, T., Ukekawa, R., … & Daub, C. O. (2010). A comprehensive survey of 3 animal miRNA modification events and a possible role for 3 adenylation in modulating miRNA targeting effectiveness. Genome research, 20(10), 1398–1410.

Burroughs, A. M., & Ando, Y. (2019). Identifying and characterizing functional 3′ nucleotide addition in the miRNA pathway. Methods, 152, 23–30.

Chang, H. T., Li, S. C., Ho, M. R., Pan, H. W., Ger, L. P., Hu, L. Y., … & Tsai, K. W.(2012, December). Comprehensive analysis of microRNAs in breast cancer. In BMC genomics (Vol. 13, No. 7, pp. 1–10). BioMed Central.

Chang, L., Zhou, G., Soufan, O., & Xia, J. (2020). miRNet 2.0: network-based visual analytics for miRNA functional analysis and systems biology. Nucleic Acids Research, 48(W1), W244–W251.

Chen, S. N., Chang, R., Lin, L. T., Chern, C. U., Tsai, H. W., Wen, Z. H., … & Tsui, K. H. (2019). MicroRNA in ovarian cancer: biology, pathogenesis, and therapeutic opportunities. International journal of environmental research and public health, 16(9), 1510.

Chiang, T. W. W., le Sage, C., Larrieu, D., Demir, M., & Jackson, S. P. CRISPR-Cas9D10A nickase-based genotypic and phenotypic screening to enhance genome editing. Sci Rep. 2016; 6: 24356.

Czabotar, P. E., Lessene, G., Strasser, A., & Adams, J. M. (2014). Control of apoptosis by the BCL-2 protein family: implications for physiology and therapy. Nature reviews Molecular cell biology, 15(1), 49–63.

D’Ambrogio, A., Gu, W., Udagawa, T., Mello, C. C., & Richter, J. D. (2012). Specific miRNA stabilization by Gld2-catalyzed monoadenylation. Cell reports, 2(6), 1537–1545.

Faehnle, C. R., Walleshauser, J., & Joshua-Tor, L. (2017). Multi-domain utilization by TUT4 and TUT7 in control of let-7 biogenesis. Nature structural & molecular biology, 24(8), 658–665.

Feng, X., Wang, Z., Fillmore, R., & Xi, Y. (2014). MiR-200, a new star miRNA in human cancer. Cancer letters, 344(2), 166–173.

Friedman, R. C., Farh, K. K. H., Burge, C. B., & Bartel, D. P. (2009). Most mammalian mRNAs are conserved targets of microRNAs. Genome research, 19(1), 92–105.

Fu, X., Meng, Z., Liang, W., Tian, Y., Wang, X., Han, W., … & Huang, W. (2014). miR-26a enhances miRNA biogenesis by targeting Lin28B and Zcchc11 to suppress tumor growth and metastasis. Oncogene, 33(34), 4296–4306.

Gutiérrez-Vázquez, C., Enright, A. J., Rodríguez-Galán, A., Pérez-García, A., Collier, P., Jones, M. R., … & Sánchez-Madrid, F. (2017). 3 Uridylation controls mature microRNA turnover during CD4 T-cell activation. Rna, 23(6), 882–891.

Heo, I., Joo, C., Cho, J., Ha, M., Han, J., & Kim, V. N. (2008). Lin28 mediates the terminal uridylation of let-7 precursor MicroRNA. Molecular cell, 32(2), 276–284.

Heo, I., Joo, C., Kim, Y. K., Ha, M., Yoon, M. J., Cho, J., … & Kim, V. N. (2009). TUT4 in concert with Lin28 suppresses microRNA biogenesis through pre-microRNA uridylation. Cell, 138(4), 696–708.

Heo, I., Ha, M., Lim, J., Yoon, M. J., Park, J. E., Kwon, S. C., … & Kim, V. N. (2012). Mono-uridylation of pre-microRNA as a key step in the biogenesis of group II let-7 microRNAs. Cell, 151(3), 521–532.

Kim, B., Ha, M., Loeff, L., Chang, H., Simanshu, D. K., Li, S., … & Kim, V. N. (2015). TUT 7 controls the fate of precursor microRNA s by using three different uridylation mechanisms. The EMBO journal, 34(13), 1801–1815.

Kim, H., Kim, J., Yu, S., Lee, Y. Y., Park, J., Choi, R. J., … & Kim, V. N. (2020). A Mechanism for microRNA arm switching regulated by uridylation. Molecular Cell, 78(6), 1224–1236.

Kim, V. N. (2005). MicroRNA biogenesis: coordinated cropping and dicing. Nature reviews Molecular cell biology, 6(5), 376–385.

Kim, Y. K., Heo, I., & Kim, V. N. (2010). Modifications of small RNAs and their associated proteins. Cell, 143(5), 703–709.

Kingston, E. R., & Bartel, D. P. (2019). Global analyses of the dynamics of mammalian microRNA metabolism. Genome research, 29(11), 1777–1790.

Kirkin, V., Joos, S., & Zörnig, M. (2004). The role of Bcl-2 family members in tumorigenesis. Biochimica et Biophysica Acta (BBA)-Molecular Cell Research, 1644(2-3), 229–249.

Knouf, E. C., Wyman, S. K., & Tewari, M. (2013). The human TUT1 nucleotidyl transferase as a global regulator of microRNA abundance. PloS one, 8(7), e69630.

Koppers-Lalic, D., Hackenberg, M., Bijnsdorp, I. V., van Eijndhoven, M. A., Sadek, P., Sie, D., … & Pegtel, D. M. (2014). Nontemplated nucleotide additions distinguish the small RNA composition in cells from exosomes. Cell reports, 8(6), 1649–1658.

Koutsaki, M., Libra, M., Spandidos, D. A., & Zaravinos, A. (2017). The miR-200 family in ovarian cancer. Oncotarget, 8(39), 66629.

Lehrbach, N. J., Armisen, J., Lightfoot, H. L., Murfitt, K. J., Bugaut, A., Balasubramanian, S., & Miska, E. A. (2009). LIN-28 and the poly (U) polymerase PUP-2 regulate let-7 microRNA processing in Caenorhabditis elegans. Nature structural & molecular biology, 16(10), 1016.

Li, J., Han, X., Wan, Y., Zhang, S., Zhao, Y., Fan, R., … & Zhou, Y. (2018). TAM 2.0: tool for MicroRNA set analysis. Nucleic acids research, 46(W1), W180–W185.

Li, S. C., Tsai, K. W., Pan, H. W., Jeng, Y. M., Ho, M. R., & Li, W. H. (2012, December). MicroRNA 3’end nucleotide modification patterns and arm selection preference in liver tissues. In BMC systems biology (Vol. 6, No. 2, pp. 1–12). BioMed Central.

Li, S. C., Liao, Y. L., Ho, M. R., Tsai, K. W., Lai, C. H., & Lin, W. C. (2012, December). miRNA arm selection and isomiR distribution in gastric cancer. In BMC genomics (Vol. 13, No. 1, pp. 1–10). BioMed Central.

Lim, J., Ha, M., Chang, H., Kwon, S. C., Simanshu, D. K., Patel, D. J., & Kim, V. N. (2014). Uridylation by TUT4 and TUT7 marks mRNA for degradation. Cell, 159(6), 1365–1376.

Lin, S., & Gregory, R. I. (2015). Identification of small molecule inhibitors of Zcchc11 TUTase activity. RNA biology, 12(8), 792–800.

Liu, J., Carmell, M. A., Rivas, F. V., Marsden, C. G., Thomson, J. M., Song, J. J., … & Hannon, G. J. (2004). Argonaute2 is the catalytic engine of mammalian RNAi. Science, 305(5689), 1437–1441.

Lu, J., Getz, G., Miska, E. A., Alvarez-Saavedra, E., Lamb, J., Peck, D., … & Golub, T. R. (2005). MicroRNA expression profiles classify human cancers. nature, 435(7043), 834–838.

Lu, P. J., Lu, Q. L., Rughetti, A., & Taylor-Papadimitriou, J. (1995). bcl-2 overexpression inhibits cell death and promotes the morphogenesis, but not tumorigenesis of human mammary epithelial cells. The Journal of cell biology, 129(5), 1363–1378.

Neilsen, C. T., Goodall, G. J., & Bracken, C. P. (2012). IsomiRs–the overlooked repertoire in the dynamic microRNAome. Trends in genetics, 28(11), 544–549.

Le Pen, J., Jiang, H., Di Domenico, T., Kneuss, E., Kosałka, J., Leung, C., … & Miska, E. A. (2018). Terminal uridylyltransferases target RNA viruses as part of the innate immune system. Nature structural & molecular biology, 25(9), 778–786.

Pfeiffer, M. J., & Schalken, J. A. (2010). Stem cell characteristics in prostate cancer cell lines. European urology, 57(2), 246–255.

Piskounova, E., Polytarchou, C., Thornton, J. E., LaPierre, R. J., Pothoulakis, C., Hagan, J. P., … & Gregory, R. I. (2011). Lin28A and Lin28B inhibit let-7 microRNA biogenesis by distinct mechanisms. Cell, 147(5), 1066–1079.

Raudvere, U., Kolberg, L., Kuzmin, I., Arak, T., Adler, P., Peterson, H., & Vilo, J. (2019). g: Profiler: a web server for functional enrichment analysis and conversions of gene lists (2019 update). Nucleic acids research, 47(W1), W191–W198.

Saiselet, M., Gacquer, D., Spinette, A., Craciun, L., Decaussin-Petrucci, M., Andry, G., … & Maenhaut, C. (2015). New global analysis of the microRNA transcriptome of primary tumors and lymph node metastases of papillary thyroid cancer. BMC genomics, 16(1), 1–18.

Song, J. J., Smith, S. K., Hannon, G. J., & Joshua-Tor, L. (2004). Crystal structure of Argonaute and its implications for RISC slicer activity. science, 305(5689), 1434–1437.

Stone, K. R., Mickey, D. D., Wunderli, H., Mickey, G. H., & Paulson, D. F. (1978). Isolation of a human prostate carcinoma cell line (DU 145). International journal of cancer, 21(3), 274–281.

Sulaiman, S. A., Ab Mutalib, N. S., & Jamal, R. (2016). miR-200c regulation of metastases in ovarian cancer: potential role in epithelial and mesenchymal transition. Frontiers in pharmacology, 7, 271.

Thornton, J. E., Chang, H. M., Piskounova, E., & Gregory, R. I. (2012). Lin28-mediated control of let-7 microRNA expression by alternative TUTases Zcchc11 (TUT4) and Zcchc6 (TUT7). RNA, 18(10), 1875–1885.

Thornton, J. E., Du, P., Jing, L., Sjekloca, L., Lin, S., Grossi, E., … & Gregory, R. I. (2014). Selective microRNA uridylation by Zcchc6 (TUT7) and Zcchc11 (TUT4). Nucleic acids research, 42(18), 11777–11791.

Ustianenko, D., Chiu, H. S., Treiber, T., Weyn-Vanhentenryck, S. M., Treiber, N., Meister, G., … & Zhang, C. (2018). LIN28 selectively modulates a subclass of let-7 microRNAs. Molecular cell, 71(2), 271–283.

Viswanathan, S. R., Daley, G. Q., & Gregory, R. I. (2008). Selective blockade of microRNA processing by Lin28. Science, 320(5872), 97–100.

Vitsios, D. M., & Enright, A. J. (2015). Chimira: analysis of small RNA sequencing data and microRNA modifications. Bioinformatics, 31(20), 3365–3367.

Wang, L., Nam, Y., Lee, A. K., Yu, C., Roth, K., Chen, C., … & Sliz, P. (2017). LIN28 zinc knuckle domain is required and sufficient to induce let-7 oligouridylation. Cell reports, 18(11), 2664–2675.

Warkocki, Z., Krawczyk, P. S., Adamska, D., Bijata, K., Garcia-Perez, J. L., & Dziembowski, A. (2018). Uridylation by TUT4/7 restricts retrotransposition of human LINE-1s. Cell, 174(6), 1537–1548.

Wyman, S. K., Knouf, E. C., Parkin, R. K., Fritz, B. R., Lin, D. W., Dennis, L. M., … & Tewari, M. (2011). Post-transcriptional generation of miRNA variants by multiple nucleotidyl transferases contributes to miRNA transcriptome complexity. Genome research, 21(9), 1450–1461.

Yang, A., Bofill-De Ros, X., Shao, T. J., Jiang, M., Li, K., Villanueva, P., … & Gu, S. (2019). 3 Uridylation confers miRNAs with non-canonical target repertoires. Molecular cell, 75(3), 511–522.

Yeom, K. H., Heo, I., Lee, J., Hohng, S., Kim, V. N., & Joo, C. (2011). Single-molecule approach to immunoprecipitated protein complexes: insights into miRNA uridylation. EMBO reports, 12(7), 690–696.

Zou, Q., Mao, Y., Hu, L., Wu, Y., & Ji, Z. (2014). miRClassify: an advanced web server for miRNA family classification and annotation. Computers in biology and medicine, 45, 157–160.

